# Characterizing the diverse cells that associate with the developing commissures of the zebrafish forebrain

**DOI:** 10.1101/2020.07.16.205153

**Authors:** J. Schnabl, M.P.H. Litz, C. Schneider, N. PenkoffLidbeck, S. Bashiruddin, M.S. Schwartz, K. Alligood, M.J.F. Barresi

## Abstract

During embryonic development of bilateral organisms, neurons send axons across the midline at specific points to connect the two halves of the nervous system with a commissure. Little is known about the cells at the midline that facilitate this tightly regulated process. We exploit the con served process of vertebrate embryonic development in the zebrafish model system to elucidate the identity of cells at the midline that may facilitate postoptic (POC) and anterior commissure (AC) development. We have discovered that three different*gfap+ astroglial* cell morphologies persist in contact with pathfinding axons throughout commissure formation. Similarly, *olig2+* progenitor cells occupy delineated portions of the postoptic and anterior commissures. These early *olig2+* progenitors demonstrate glial-like morphologies despite the lack of a myelination marker. Moreover, we conclude that both the *gfap+* and *olig2+*progenitor cells give rise to neuronal populations in both the telencephalon and diencephalon. Interestingly, these varied cell populations showed significant developmental heterochrony between the telencephalon and diencephalon. Lastly, we also showed that *fli1a+* mesenchymal cells migrate along the presumptive commissure regions before and during midline axon crossing. Furthermore, following commissure maturation, specific blood vessels formed at the midline of the POC and immediately ventral and parallel to the AC. This comprehensive account of the cellular populations that correlate with the timing and position of commissural axon pathfinding has supported the conceptual modeling and identification of the early forebrain architecture that may be necessary for proper commissure development.

## 1 Introduction

Development of the central nervous system (CNS) is a complex process that requires the coordinated generation of the correct number, position and differentiation of neuronal and glial lineages. Vertebrate brain development in particular involves the creation of a diverse array of cell types, formation of complex and interacting neural circuits, and multidimensional cell migration with tissue level morphogenesis. The relatively simple vertebrate model system of the zebrafish, *Danio rerio*, has provided a particularly tractable system to probe the mechanisms of brain development because it offers direct and transparent access to the early developing embryo with the powerful advantages of genetic manipulation (Pathak and Barresi, 2020).

The commissure is one of the most striking structures in the CNS that is characteristic of all bilaterally symmetrical organisms. During embryonic development pathfinding axons are guided to and across the midline forming a commissure at stereotypical locations along the anterior to posterior axis of the organism. Commissures are a defining architecture of the CNS, and understanding the regulation of their position and growth remains a fundamental pursuit in the field of developmental neurobiology (Hjorth and Key, 2004; Kaprielian, Runko, and Imondi, 2001).

The first two commissures to form in zebrafish appear in the forebrain, constituting the post optic commissure (POC) in the diencephalon and the anterior commissure in the telencephalon (Wilson and Easter, 1991; Chitnis and Kuwada, 1990). The first pioneering axons to cross the forebrain midline and mediate subsequent commissural crossing events do so in contact with glial cells which express the astroglial marker Glial fibrillary acidic protein (Gfap). These Gfap+ glial cells are arranged in a bridge-like pattern which subsequent commissural axons navigate across (Barresi et al., 2005). Prior to commissure formation, processes of these Gfap+ cells appear to ubiquitously cover the basal or superficial most extents of the forebrain, however Gfap labeling soon condenses into midline spanning swaths of astroglia corresponding to the presumptive diencephalic and telencephalic commissural regions (Barresi et al., 2005). Moreover, axonal growth is quantitatively correlated with the movement of these astroglial cells over the course of commissure formation, strongly suggesting that these “glial bridges” provide a supportive substrate for pathfinding commissural axons (Barresi et al., 2005; Schwartz et al., 2020).

Historically, the study of commissure development has largely focused on the behavior of axon pathfinding, which has led to a vast and important literature on the role of molecular axon guidance cues (Kaprielian, Runko, and Imondi, 2001; Zou and Lyuksyutova, 2007; Fothergill et al., 2014; Tosa et al., 2015; Friocourt and Chédotal, 2017). Despite recent advances and increased power of genetic techniques and microscopy, less attention has been paid to the cellular and matrix microenvironments that pathfinding commissural axons use to pioneer the midline during development (Stanic et al., 2016; Oliver et al., 2019). However, concomitant with the midline crossing events of commissural axons, is the development and expansion of a seemingly non-deterministic number of other cells with their own currently undefined array of dynamic behaviors. How these other cell types may contribute to forebrain development, and more specifically, how they may influence commissure formation has been a neglected question. It is well understood that during early neural tube development, neuroepithelial cells are the initial progenitors for primary neurons, glia, and more derived progenitor cell populations (H. Kim et al., 2008; Ravanelli and Appel, 2015; Zhou, Wang, and Anderson, 2000; Garcia-Lopez and Martinez, 2010; Kriegstein and Alvarez-Buylla, 2009a; De Juan Romero and Borrell, 2015). During initial neurogenesis, many neuroepithelial cells transition into radial glial cells (RGC) and retain their stem cell functions throughout the life of a zebrafish (Malatesta, Hartfuss, and Götz, 2000; Noctor et al., 2001; Miyata et al., 2001; Alvarez-Buylla, Garcia-Verdugo, and Tramontin, 2001; Kriegstein and Alvarez-Buylla, 2009b; Adolf et al., 2006; Pinto and Götz, 2007; Chapouton et al., 2006). Importantly, RGCs are established throughout the CNS well before any axonogenesis has occurred (Bernhardt, 1999; Ross, Parrett, and S. Easter, 1992; Johnson et al., 2016; De Juan Romero and Borrell, 2015; Malatesta, Hartfuss, and Götz, 2000), and they have been found to be essential for both the architectural integrity of the CNS and the generation of neuronal and glial populations (Johnson et al., 2016; Smith et al., 2014; Casper and McCarthy, 2006). The characteristic radial glial morphology exhibits a prominent basal process that extends from the cell body in the ventricular zone to terminating at the pial surface with a large membranous structure known as an endfoot (Alberto Docampo-Seara et al., 2019; Chanas-Sacre et al., 2000; Lindsey et al., 2018; H. Kim et al., 2008; Götz and Barde, 2005). The basal process and its terminating endfoot are often marked by the intermediate filament Gfap (Alberto Docampo-Seara et al., 2019; Lindsey et al., 2018; H. Kim et al., 2008; Götz and Barde, 2005; Marcus and Easter Jr, 1995). In both mouse and zebrafish, pathfinding axons have been shown to come in direct contact with these Gfap+ radial glial fibers during commissure development (Marcus and Stephen S Easter, 1995; Stier and Schlosshauer, 1998; Eng, Ghirnikar, and Lee, 2000; Nielsen and Jørgensen, 2003).

In the mammalian forebrain, populations of astroglia, mostly radial glia, have been suggested to form cellular substrates that support the growth of cerebral commissures (Silver et al., 1982; Shu and Richards, 2001; Shu, Puche, and Richards, 2003; Mason and Sretavan, 1997) as well as the optic chiasm, which is pioneered by axons of retinal ganglion cells from the retina (Marcus and Stephen S Easter, 1995; Petros, Rebsam, and Mason, 2008). Commissural axons migrate medially towards the midline through the dense field of radial fibers (Silver et al., 1982). In the mouse, discrete populations known as the glial sling and glial palisade exist at the forebrain midline, while the induseum griseum and the glial wedge are positioned in more peripheral locations. It has been proposed that the combination of these different astroglial populations serve to provide both permissive substrates at the midline as well as secreted non-permissive cues from the peripheral regions. Together glial cells help to channel commissural and retinal axons across the midline (Shu and Richards, 2001; Marcus and Stephen S Easter, 1995; Petros, Rebsam, and Mason, 2008). Although, little documentation currently exists to support the existence of astroglial subtypes in zebrafish, the *no-isthmus (noi; pax2a)* mutants have a loss of reticulate astrocytes in the glia limitans at the optic nerve, which results in pathfinding defects and impaired fasciculation by axons of the optic chiasm and POC (Macdonald et al., 1997). As alluded to previously, Gfap+ astroglial fibers develop into condensed midline bridges in the diencephalon and telencephalon, and directly interact with commissural and retinal axons during and after midline crossing (Barresi et al., 2005; Schwartz et al., 2020).

Astroglial cells are not the only cells to associate with commissural axons. Oligodendrocytes ultimately form dense associations with axons in order to myelinate the tracts and nerves of the CNS (Alghamdi and Fern, 2015; Fontenas and Kucenas, 2018; Lyons and Talbot, 2015; Ackerman and Monk, 2016). While mature oligodendrocytes are not present in the zebrafish CNS until after 3 days postfertilization (48 hours after the first POC axons cross the diencephalic midline), olig2+ oligodendrocyte progenitor cells (OPC) are present in the forebrain throughout commissure formation (Lyons and Talbot, 2015; Ackerman and Monk, 2016; Yoshida and Macklin, 2005). How these known neurogenic and gliogenic progenitor cells may be influencing early commissure development remains a mystery. Furthermore, during the period of commissure formation, cranial neural crest cells migrate at the periphery of the CNS, where they will eventually contribute to the development of sensory ganglia and head cartilage (Barrallo-Gimeno et al., 2004; Wada et al., 2005; Couly and Le Douarin, 1987; Graham, Begbie, and McGonnell, 2004; Le Douarin et al., 2004). Lastly, endothelial cell intrusion into the CNS is required for cerebral vasculogenesis (Lawson and Weinstein, 2002; Jin et al., 2005; Isogai, Horiguchi, and Weinstein, 2001). The spatial and temporal coordination of commissure construction with OPC maturation, cranial neural crest development and blood vessel formation has never been comparatively characterized.

In this study, we took a broad perspective in order to shed light on the diversity of cell types that may be contributing to the guidance of commissural axons across the midline of the zebrafish forebrain. We show that several morphologically distinct cells of the astroglial lineage associate with pathfinding commissural axons, and that these glial cells appear to express different radial fiber antigens which are expressed in both overlapping and differential patterns. In addition, we document the presence of *olig2:EGPF+*progenitor cells for neuronal lineages in the early forebrain between the developing AC and POC. Despite the lack of myelin markers at these early stages, the membranous processes of these OPCs were seen intimately associated with POC axons. Soon after commissure formation, we also identified the emergence of serotonergic progenitor cells within the tracts of the forebrain commissures and surrounding areas. Notably, both the serotonergic and OPCs populations of the telencephalon exhibited developmental heterochrony with these same populations in the diencephalon. In both cases, despite the earlier incidence of commissure formation, oligodendroglial, and neuronal populations in the diencephalon, development and expansion of these populations in the telencephalon rapidly surpasses that found in the diencephalon. Finally, our characterization of the timing of endothelial cell invasion and vascular development over the course of forebrain commissure formation revealed an early population of mesenchymal cells that prefigure the commissural regions. Additionally, endothelial cells temporally and spatially mirrored the growing AC, while only a midline restricted vessel was observed growing into the diencephalon after POC formation. In this study, we document the concurrent development of progenitor, glial, neuronal, and vascular cells with pathfinding commissural axons of the zebrafish forebrain. This characterization has highlighted the complex yet probable importance these varied cell types play in the guidance and maintenance of commissures and calls for further investigations into the functional roles these cell types may play in coordinating and contributing to forebrain development.

## 2. Materials and Methods

### Wild-type and transgenic fish lines

Fish lines were maintained in the Smith College Animal Quarters according to Smith College Institutional Animal Care and Use Committee (IACUC) and AAALAC regulations. Wild type embryos were from the AB strain provided by C. Lawrence (Harvard University) and Tübingen strain from ZIRC (Eugene, OR). Axons were visualized with the *tg(3.6f gap43: GFP)* trans genic line driven by a 3.6kb regulatory region of the axon growth-associated gene *gap43* (Udvadia, 2008). *tg(olig2:GFP)* and *tg(olig2:DsRed)*lines driven by 1.8 kb of the *olig2* regulatory region, were provided by Bruce Appel (Shin et al., 2003). Glial membrane reporter lines *tg(gfap:GFP-CAAX)* and *tg(gfap:mCherry-CAAX)* were designed using the promoter originally designed for the *tg(gfap:GFP)* line (Bernardos and Raymond, 2006). We used the *tg(HuC:mCherry)* line driven by 4.6 kb of upstream elements of the *elavl3/HuC* gene (Park et al., 2000) to examine the distribution of neuronal populations. The *tg(fli1a:EGFP* line was used to examine the localization of neural crest and vascular contributions to the forebrain (Lawson and Weinstein, 2002). Examination of vascular contributions to the forebrain without neural crest contributions was done using the *tg(flk1a(vegfr):EGFP)* (Jin et al., 2005). We visualized clonal populations of radial glia using the *tg(ubi:zebrabow M)* and 7.4 kb *gfap* promoter *tg(Cre^ERT2^)* lines, as described below (Bernardos and Raymond, 2006).

### Whole Mount Immunocytochemistry

Embryos were grown at 23.5-28.5°C and staged in hours post-fertilization (hpf) (Kimmel, Ballard, et al., 1995). Immunoctyochemistry was carried out as described previously (Johnson et al 2014; Johnson et al, 2016; Schwartz et al., 2020). Briefly, embryos were fixed with 4% paraformaldehyde with 10% dimethylsulfoxide (DMSO) for 3 hrs at room temp or 1 hour at room temp followed by 16 hours at 4° C. Washed embryos were dehydrated in a Methanol series that culminated with a −20°C incubation in 100% acetone for 4 minutes or 7 minutes (for ages 30 hpf or older). Following rehydration in phosphate buffered saline with 2% v/v Triton x-100 (PBS-Tx), 36 and 48 hpf embryos were also treated with 0.5% collagenase for 30 minutes. Embryos were then blocked for non-specific antibody binding for 1 hour at room temperature in a solution of PBS-Tx with 5% w/v bovine serum albumin fraction V, 1% v/v DMSO, and 10% v/v normal goat serum. Blocked embryos were incubated in primary antibody diluted in blocking solution at room temp for 2 hours or at 4° C for 16 hours. Following washes with PBS-Tx embryos were blocked again for 30-60 minutes prior to incubation in secondary antibody solution for 2 hours at room temperature or at 4° C for 16 hours. Labeled embryos were maintained in 70% glycerol at 4° C prior to imaging.

### Antibodies

Primary antibodies used include mouse IgG2B anti-Acetylated tubulin (Sigma, T6793, 1:800), mouse IgG1 anti-Zrf-1 (ZIRC, Zrf1, 1:20), mouse IgG1 anti-Zrf-2 (ZIRC, Zrf2, 1:100), mouse IgG1 anti-Zrf-3 (ZIRC, Zrf3, 1:200), mouse IgG1 anti-Zrf-4 (ZIRC, Zrf4, 1:100), chicken anti-mCherry (Millipore, AB356481, 1:1000), rabbit anti-GFP (Invitrogen A-11122 1:600), mouse IgG2b anti-HuC/D (Invitrogen, A21271, 1:250), and mouse IgG1 anti-pH3 (Cell Signaling Technology, 9706S, 1:500). Secondary antibodies used include goat anti-rabbit 488 (Invitrogen, 1:200), goat anti-mouse Igg1 647 (Invitrogen, 1:200), goat anti-mouse IgG1 488 (Invitrogen, 1:200), goat anti-mouse IgG1 594 (Invitrogen, 1:200), goat anti-mouse IgG2b Alexa 647 (Invitrogen, 1:200) goat anti-mouse IgG2b 594 (Invitrogen, 1:200), goat anti-mouse IgG2b 488 (Invitrogen, 1:200), and goat anti-chicken 594 (Invitrogen, 1:200).

### Confocal Microscopy

The anterior head regions of immunolabeled embryos were dissected off at the level of midbrain just posterior to the eyes and mounted in 70% glycerol with the POC region oriented closest to the glass cover slip. With the exception of the data presented in figures 4 and 8, all samples were imaged on a Leica SP5 laser scanning confocal microscope (LSCM) with Leica HC apochromat (CS2) 63X oil objective (0FN25/E) with a numerical aperture of 1.4 and a 1.5 optical zoom was used during acquisition. Samples used to support the data in figures 4 and 8 were acquired with Zeiss’s AxioImager Z1 with Apotome, and Z-stacks were collected with an optical sectioning thickness of 0.53 *μ*m at 40X and processed using Zen software (Zeiss). Samples collected on the LSCM were imaged at a pixel resolution of 1024 by 1024 with an additional 4-line averaging. Z-stacks of the forebrain were collected for each mounted tissue at an optical step size of 0.21 *μ*m, however when photobleaching was detected a step size of 0.38 *μ*m. The resulting Leica (lif) files were processed using Fiji to make projections and z-slice movies Sbalzarini, 2016; Schindelin et al., 2008. Additional 3D rendered projections and movies were made with Amira software, using Volren settings. The levels of channel intensity (between 0 and 255) and opacity (0-1) were adjusted to enable maximal visualization of each channel without saturation, with the lower limit of intensity set between 5-10, and upper limit between 125 and 150 Stalling, Westerhoff, and Hege, 2005 The opacity for non-AT channels was set to between .5 and .75 to enable visualization of the AT channel.

### Gastrula Stage Transplants

Gastrula-staged cell transplantation was carried out as described in (Deschene and M. J. Barresi, 2009). Briefly, donor transgenic embryos and wild-type host embryos were hand dechorionated at 4 hpf and placed into a 2% agarose transplant well plate filled with antibiotic embryo rearing medium (15 mM NaCl, 0.5 mM KCl, 1.0 mM MgSO4, 150 *μ*m KH2PO4, 1.0 mM CaCl2, 0.7 mM NaHCO3, 0.5 mg/L Methylene Blue). Using a TransferMan NK2 microinjector (Eppendorf) donor cells were extracted from the *tg(gfap:GFP)*^mi2002^ or *tg(olig2:EGFP)*^vu12^ transgenic gastrula halfway from the shield to the animal pole and transplanted into the same region of 6 hpf wild-type host embryos. These embryos were raised to 30 hpf at 28.5° C in antibiotic embryo medium and then fixed and processed for whole-mount immunocytochemistry.

### Zebrabow Cre^ERT2^ Induction

We generated small populations of fluorescent astroglial cells throughout the zebrafish nervous system using a *tg(*ubi:Zebrabow M) and *tg(gfap:cre^ERT2^)* double transgenic line. Embryos were first manually decorated at 4 hpf and then treated with 4-hydroxytamoxiphen (Sigma cat 94873) between 6-8 hpf in the absence of light. Embryos were fixed based upon protocol at 28 hpf embryos and then processed for whole-mount immunocytochemistry, which included immunolabeling with the rabbit anti:GFP antibody and with imaged the exclusion of the 594 nm (cherry) channel.

### ΔScope Data Processing

The quantitative distribution of the anti-Zrf immunolabeling was analyzed with ΔScope as described in Schwartz et al. 2020. In brief, following imaging on the confocal microscope, data from each anti-Zrf labeled sample was first cropped to focus on the POC and the background signal eliminated, after which the image converted to an HDF5 (.h5) file type using the HDF5 plugin with Fiji (The HDF Group, 1997). AT and Zrf1-4 channels were then processed using ilastik to improve the signal to noise ratios and generate signal probability values for each pixel. Output ilastik files for Zrf1-4 experimental groups were finally processed using the ΔScope workflow identical to (Schwartz et al., 2020).

## 3 Results

This investigation attempted to provide a more comprehensive characterization of the cellular context in which commissural axons grow during embryonic forebrain development. The first commissural axons of the forebrain navigate across the midline during an early stage of brain development that is particularly dynamic. During this developmental period, many cell behaviors are occurring simultaneously as the commissure is forming, which includes progenitor cell division at the ventricular zones, various types of cell migration, epithelial morphogenesis, and cell differentiation (Kriegstein and Alvarez-Buylla, 2009b; Lewis and Eisen, 2003; Lindsey et al., 2018). Documentation of the timing and spatial relationship between pathfinding commissural axons and these varied transient and permanent cell types in the forebrain has not been completed. This study sought to establish a base-line understanding of the varied cell types that may provide important axon guidance to commissural neurons. We take advantage of the zebrafish model system’s accessibility for high resolution microscopic analysis and relatively simple embryonic brain architecture as compared to its mammalian vertebrate counterparts (Pathak and Barresi, 2020).

### 3.1 Defining the Differentiated Forebrain

To identify cell types that may be influencing commissure formation, we first sought to define the most prominent cells residing in commissural regions at a stage of mature commissural anatomy. The 48 hour post fertilization (hpf) forebrain represents a developmental stage where the optic chiasm, post optic, and anterior commissures are all robustly fasciculated. For this characterization, we targeted the four most well-known classes of cells in the developing brain, progenitor cells, neurons, glial cells, and vasculature. Throughout this study we took advantage of commercial antibodies that recognize Acetylated Tubulin (AT) to broadly label all the axons of the forebrain. Immunocytochemistry was performed in conjunction with fluorescent transgenic reporter lines specific for each of the aforementioned different classes of cells *tg(gfap:GFP-CAAX)*, radial glial cells (Johnson et al., 2016); *tg(huC:mCherry)*, pan-neuronal (Park et al., 2000); *tg(olig2:EGFP)*, oligodendroglial (Shin et al., 2003); *tg(fli1a:EGFP)*, endothelial cells (Lawson and Weinstein, 2002).

As previously reported, AT labeling marked the AC, that spans the telencephalic midline, and the POC and optic chiasm, that span the diencephalic midline (Wilson and Easter, 1991); Figure 1 A,B). At this late stage, the density of axons which form the optic chiasm can obscure significant portions of the underlying POC axons (Figure 1 A,L,V,GG, blue; see also Sup. Mov. 1, blue). It is known from a collection of studies that the neurons pioneering the AC and POC originate in part from the dorsal and ventral rostral clusters, respectively (Hjorth and Key, 2004; Bak and Fraser, 2003).

**Figure 1:**
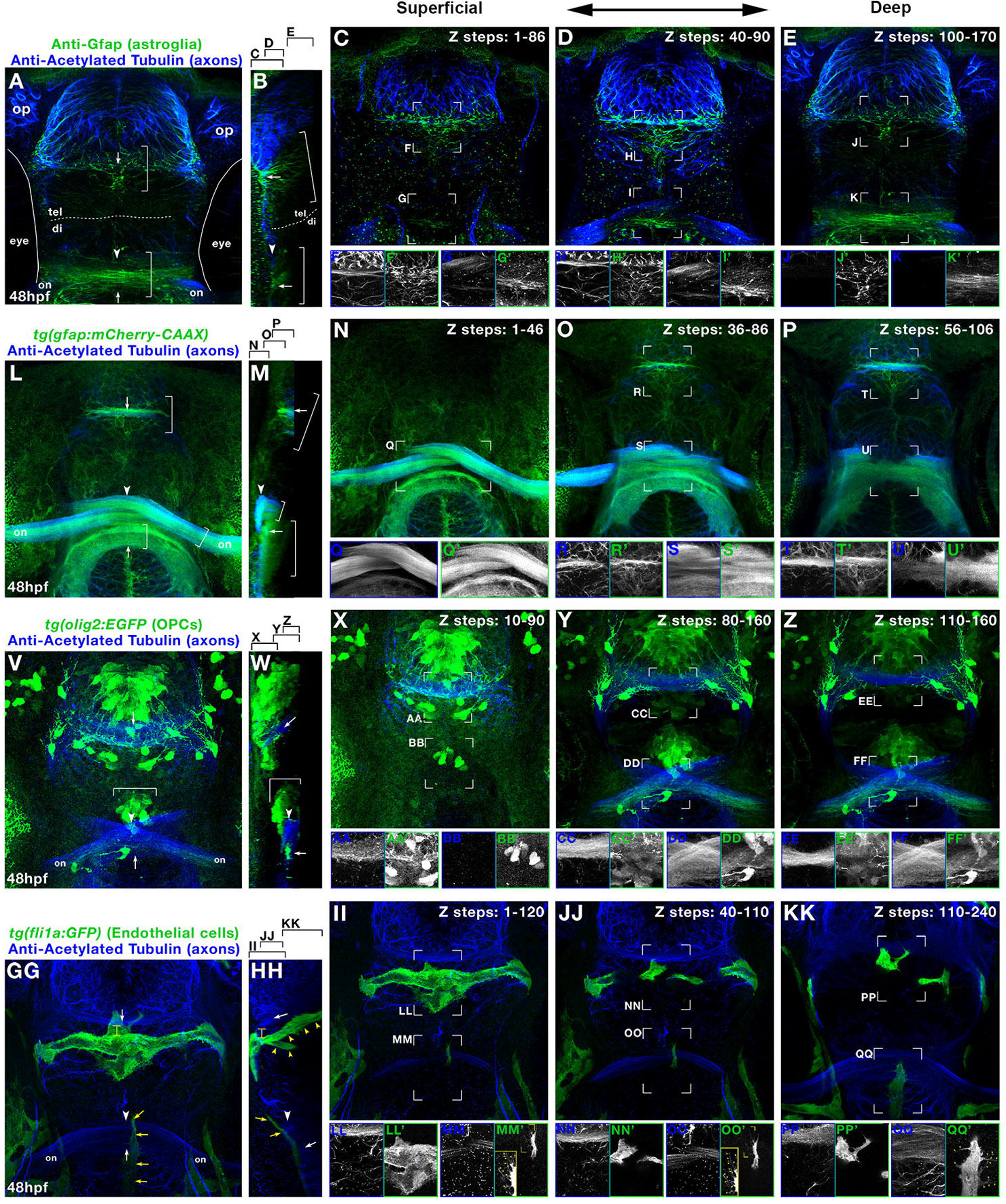
The mature embryonic zebrafish forebrain is composed of commissural axons, neurons, *fli1a+* endothelial cells, and *olig2+* cells. Color composites of anti-Acetylated Tubulin (anti-AT, blue in all) and Anti-Gfap (A-E, green), *gfap:cherry-caax* (L-P, green), *olig2:EGFP* (V-Z, green), *fli1a:EGFP* (GG-KK, green). A,L,V,GG) MIP color composites, with respective sagittal sections (A,L,V,GG). C-E, N-P, X-Z, II-KK) Selected Z slices from (A, L, V, GG), respectively, from more anterior (C, N, X, II) to more posterior (E, P, Z, KK). Monochrome insets of signal from composite z-slice selections: (F-J,Q-T,AA-EE,LL-QQ) anti-AT, (F’-J’) anti-Gfap, (Q’-T’) *gfap:cherry-caax*, (AA’-EE’) *olig2:EGFP*, (LL’-QQ’) *fli1a:EGFP*. MM’, OO’) Yellow boxes with zoom of yellow bracketed regions. Markings: (yellow arrows) endothelial cell processes, (white arrowheads) optic chasm, (top white arrow) anterior commissure, (bottom white arrow) post optic commissure. Abbreviations: op, olfactory pit; on, optic nerve; tel, telencephalon; di, diencephalon. Z-step = 0.21*μ*m. hpf, hours post fertilization

We have previously shown that the forebrain commissures develop in direct contact with astroglial cells expressing Gfap (Figure 1 A, green; (Barresi et al., 2005; Schwartz et al., 2020)). At 48 hpf, Gfap densely populated the most superficial/basal regions of the forebrain and specifically overlapped with and interlaced through each commissure (Figure 1 A-E, brackets; Sup. Mov. 1 A). These results are consistent with previous studies showing Gfap in the endfeet of radial glial cell processes anchored to the basal surface (Sarthy, Fu, and Huang, 1989). Using the membrane-labeling astroglial transgenic line *tg(gfap:GFP-CAAX)* (Johnson et al., 2016), we show here that in both the telencephalon and diencephalon, astroglial cell membranes completely overlapped the axons of the AC and POC respectively (Figure 1L-P, brackets; Sup. Mov. 1 B). The prominent correlation of *gfap:GFP-CAAX* with Gfap protein and commissural axons is consistent with the subcellular localization of the Gfap intermediate filaments to the more basal astroglial cell processes (Figure 1 A,B vs L,M). Interestingly, *gfap:GFP-CAAX* also revealed deep expression that extended into and significantly overlapped with the optic nerves (Figure 1 L,M, small bracket). Our results suggest that in the 48 hpf zebrafish forebrain, Gfap-rich radial glial cell processes comprise a significant portion of the commissural regions.

Another essential class of cells in the CNS is oligodendroglia, which principally function to provide the insulating myelin for axonal tracts and commissures (Alghamdi and Fern, 2015; Borodovsky et al., 2009; Lewis and Eisen, 2003; Ackerman and Monk, 2016). True myelinating oligodendrocytes most-commonly identified by expression of myelin basic protein (MBP/*mbpa*) are not present in the CNS before 5 days post fertilization (dpf) in zebrafish (Pogoda et al., 2006); however, *olig2+* oligodendrocyte progenitor cells (OPCs) are present for much of the late embryonic and larval periods of CNS development (Ackerman and Monk, 2016; Lyons and Talbot, 2015; Mitew et al., 2014; Ravanelli and Appel, 2015; Shin et al., 2003). Taking advantage of the *tg(olig2:EGFP)* transgenic line, we investigated the spatial relationship that *olig2+* OPCs have with the forebrain commissures at 48 hpf. We confirm that *olig2+* OPCs are present throughout the forebrain (Borodovsky et al., 2009), yet we show here significant differences in OPC positioning in the telencephalon and diencephalon (Figure 1 V-Z; Sup. Mov. 1 C). We observed that *olig2+* OPCs densely populated the telencephalon, whereas a significantly smaller population of OPCs were present in the diencephalon (Figure 1 V,W, bracket). Moreover, with the exception of a relatively smaller population of more ventrally located cells, the *olig2+* OPCs in the diencephalon were focused into a relatively small cluster of cells positioned directly on the midline just dorsal to the optic chiasm (Figure 1 V-Z, bracket and boxed areas). Interestingly, despite the lack of known MBP expression at 2 dpf, some *olig2+* OPCs had multiple long cellular processes that both made direct contact with the tracts of the commissures, and was enmeshed within the AT signal of the commissures (Figure 1 X-Z). Notably, the two isolated cells ventral to the diencephalic *olig2+*cell cluster extended processes into both the POC and optic nerves (Figure 1 V,W, arrow). The observed interactions between AT labeled axons and *olig2+*progenitor cell processes suggest that these OPCs are demonstrating pre-myelinating behaviors and/or developing into neurons whose axons fasciculate with the commissures.

Brain development is also influenced by interactions with cells that originate externally to the ectoderm. Although first derived from the neural ectoderm, neural crest cells are arguably the most prominent embryonic cell type external to the CNS that is well known to interact with the brain, as they contribute to the development of peripheral sensory ganglia and the craniofacial skeleton (Kimmel, Miller, and Keynes, 2001; Etchevers et al., 1999; Creuzet, 2009; Le Douarin et al., 2004; Graham, Begbie, and McGonnell, 2004). Additionally, development of the blood-brain barrier requires the infiltration of migrating vascular cells that form endothelial-cell lined blood vessels throughout the brain (Lawson and Weinstein, 2002; Gross and Hanken, 2008). Therefore, it was important for our characterization to include the spatial distribution of developing vascular and cartilaginous structures in the proximity of the forebrain commissures. To visualize these cell populations, we took advantage the *tg(fli1a:EGFP)* transgenic line which labels both populations (Lawson and Weinstein, 2002). At 48 hpf, we observed discrete populations of *fli1a:EGFP+* cells in the regions of both the telencephalon and diencephalon that resembled endothelial cells undergoing vasculogenesis (Figure 1 GG-KK; Sup. Mov. 1 D). More specifically, the palatocerebral artery (PLA) (Isogai, Horiguchi, and Weinstein, 2001) was seen forming in the telencephalon in a position parallel to but ventrally displaced from the AC (Figure 1 GG, HH, yellow bracket). At 48 hpf we also captured the initiation of deeper endothelial cell sprouts behind the AC growing in a dorsoventral direction at the midline in the telencephalon (Figure 1 HH, yellow arrowheads). In contrast, vasculogenesis in the diencephalon at 48 hpf was restricted to a single stream of dorsally-growing endothelial cells along the midline, found at the extreme basal-most extent of the brain, superficial to even the POC (Figure 1 GG,HH, yellow arrows). Each of these structures is consistent with previously described patterns of early vascularization of the forebrain (Lawson and Weinstein, 2002), however, by placing them in the context of the established commissures, this study has provided a better understanding of vascular position and potential vessel-axon interactions. Importantly, we did not observe any *fli1a:EGFP+* labeling suggestive of early cartilaginous structures like the mandibular arches, cranial vault or frontal nasal mass that are known to be present at 56 hpf (Lawson and Weinstein, 2002; Creuzet, 2009). Our initial characterization of the diversity of tissue types that interact with the mature commissures of the forebrain has identified progenitor, neuronal, glial and vascular cells that may all play important roles in shaping the earlier development of these commissures. We next extended this characterization to evaluate the specific contributions these varied cell types have to the forebrain before and during commissural axon pathfinding across the midline.

### 3.2 Commissural Axon - Progenitor Cell Interactions

The first commissural axons to cross the midline of the diencephalon and telencephalon do so during a period of intense progenitor cell division and differentiation (Bak and Fraser, 2003; Ross, Parrett, and S. Easter, 1992; Schier, 2001; Rallu, Corbin, and Fishell, 2002; Schuurmans and Guillemot, 2002; Lindsey et al., 2018; Beretta et al., 2017). Therefore, we propose that exploring the origin and spatial relationship that the varied forebrain progenitor cell populations have with pathfinding axons may shed crucial light on how these predominant stem cell populations may influence commissure formation. The early CNS first develops from the ectoderm as a sheet of neuroepithelial cells undergo several morphogenetic events to form the neural tube (H. Kim et al., 2008; Lewis and Eisen, 2003; Götz and Barde, 2005). These early neuroepithelial cells are understood to be the first population of progenitor cells fueling primary neurogenesis and the birth of new stem cell populations, namely radial glial cells and Olig2 progenitor cells (H. Kim et al., 2008; Götz and Barde, 2005; De Juan Romero and Borrell, 2015; Lyons and Talbot, 2015; Pringle et al., 1996; H. Kim et al., 2008). Although markers which are specific for neuroepithelial cells in zebrafish do not exist, radial glial cells appear to be predominantly responsible for the majority of progenitor cell development in the CNS soon after gastrulation, and thus constitute the most abundant stem cell present during commissure formation (Johnson et al., 2016).

### 3.3 Radial Glial Development

We have previously identified Gfap+ cell processes to be present before and throughout forebrain commissure formation in zebrafish (Barresi et al., 2005). In addition, by using ΔSCOPE, a new computational method for 3-Dimensional image quantification, we have recently described the concordant behavior of Gfap-labeled fibers and POC axons during midline crossing. More specifically, as POC axons pathfind over the midline and remodel into a commissure, the Gfap+ glial cell fibers become increasingly condensed around the commissure overtime much as observed in (Figure 1 A-K; (Schwartz et al., 2020)). However, the identity of these commissure-associated cells is particularly uncertain. We hypothesized that Gfap+ astroglia in the commissure region represent a heterogeneous population of progenitor cells as well as structural cell types of astroglial origin. To explore this hypothesis, we conducted three experimental approaches: First, we characterized the temporal and spatial distribution of astroglial cell markers over the timecourse of commissure formation. Second, we performed gastrula-staged cell transplantation procedures with glial-labeling transgenic lines to visualize the position and morphology of isolated cells. Lastly, we confirmed these cells were of astroglial origin by using the *tg(gfap:CRE^ERT2^;ubi:zebrabow)* transgenic line to clonally label gfap+ cells in the forebrain.

Gfap is a very well characterized and established marker of astroglial cells (H. Kim et al., 2008; Johnson et al., 2016; Eng, Ghirnikar, and Lee, 2000; Götz and Barde, 2005; Docampo-Seara et al., 2018; Lam, März, and Strähle, 2009; Kriegstein and Alvarez-Buylla, 2009a; R. L. Bernardos and P. A. Raymond, 2006)); therefore, we first attempted to characterize the position of whole Gfap-expressing cells associated with forebrain commissures. We immunolabeled embryos from the membrane-marking astroglial transgenic reporter line *tg(gfap:GFP-CAAX)* at 22 hpf, 24 hpf, 28 hpf and 36 hpf for both AT (axons) and Gfap (astroglial intermediate filaments). We observed that just prior to and well after midline crossing of POC axons, *tg(gfap:GFP-CAAX)*-marked cells showed refined localization to midline-spanning “bridges” in both the telencephalon and diencephalon (Figure 2 A,E,I,M (Magenta), Fig 2A-D). Moreover, Gfap intermediate filament protein expression is prominently seen within the *tg(gfap:GFP-CAAX)*-marked cell processes, and was similarly refined into midline-spanning bridges throughout the timecourse (Figure 2; Green and Magenta respectively; see also Sup. Mov. 2 A-D); (Barresi et al., 2005; Schwartz et al., 2020)). By 28 hpf, both Gfap and astroglial cell membranes were densely packed around the commissures and infiltrating in between fasciculated axons as revealed in Y-Z axis rotations of these labeled forebrains (Figure 2 B-D, F-H, J-L, N-P). These results suggest that pathfinding commissural axons grow across the midline through a meshwork of astroglial cells. This strong association of commissural axons with Gfap+ cells and fibers continued through 36hpf (Figure 2 M-P) and into 48 hpf (see Figure 1), which suggests that these astroglial cells play an active role in both the establishment and further development of the forebrain commissures.

**Figure 2:**
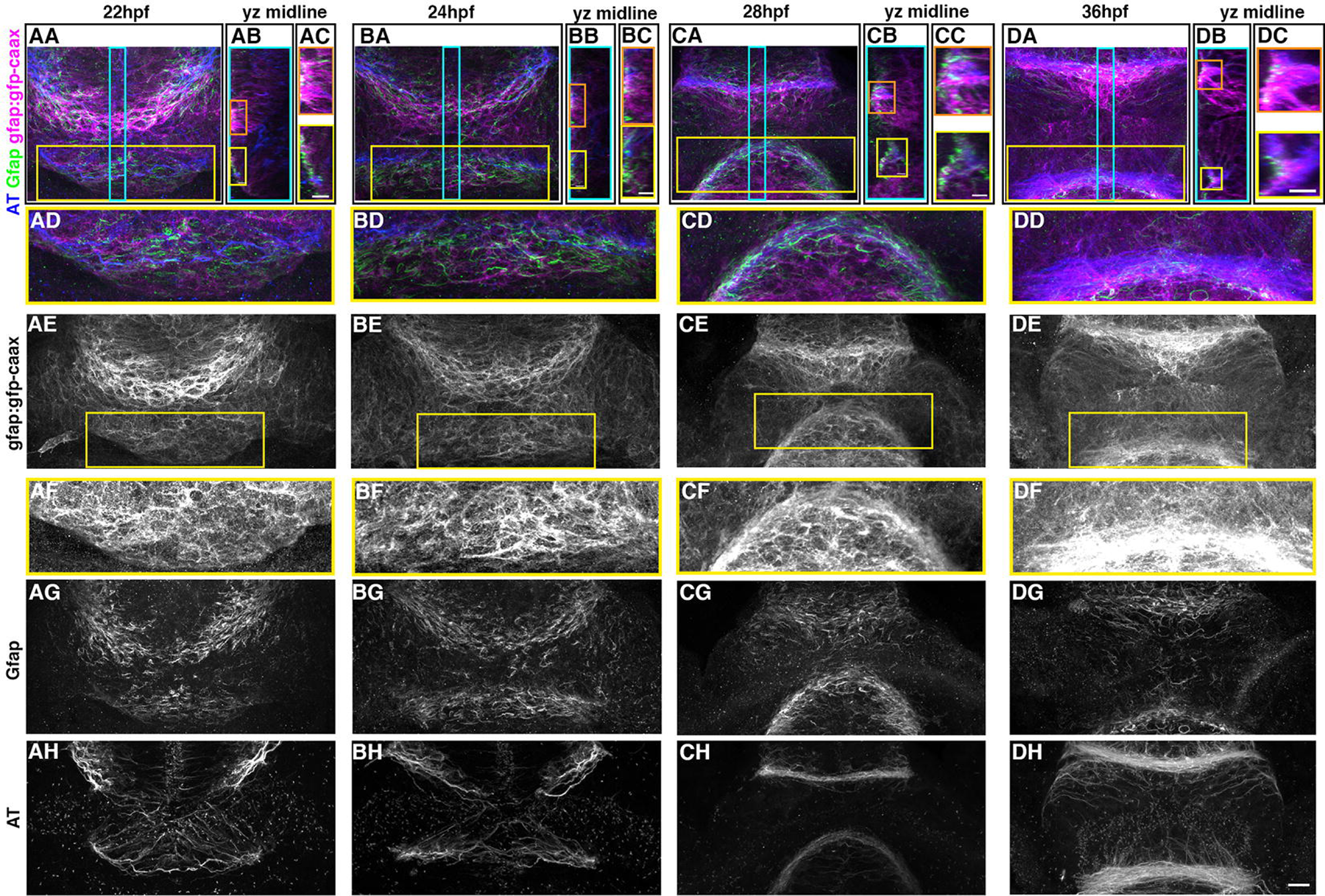
Gfap+ glial bridges span the telencephalon and diencephalon, and develop concomitantly with the post optic commissure and anterior commissure. AA, BA, CA, DA) frontal view color composites with *tg(gfap:gfp-caax)* (magenta), immunolabeled anti-Zrf1 (Gfap, green) and anti-Acetylated Tubulin (AT, blue) at 22, 24, 28 and 36 hpf, respectively (scale bar = 11*μ*m). AB, BB, CB, DB) Sagittal midline sections shown in turquoise box in (AA, BA, CA, DA; scale bar = 6*μ*m), with insets (AC, BC, CC, DC) of the anterior commissure (orange boxes; scale bar = 6*μ*m) and post optic commissure (yellow boxes; scale bar = 6*μ*m). AD, BD, CD, DD) Insets from yellow boxed frontal view in (AA, BA, CA, DA) highlighting the POC and diencephalic glial bridge. Monochrome channels of (AA,BA,CA,DA), respectively, are: (AE, BE, CE, DE) *tg(gfap:gfp-caax)*, (AG, BG, CG, DG) Gfap, (AH, BH, CH, DH) AT. AF, BF, CF, DF) Insets of (AE, BE, CE, DE) marked by yellow boxes, with uniformly increased intensity levels to show glial membranes of the diencephalic glial bridge. hpf, hours post fertilization

In both chick and explant cultures, axons have been observed to migrate preferentially along glial cell processes as opposed to the cell’s soma (Stier and Schlosshauer, 1998; Stier and Schlosshauer, 1995). To more comprehensively examine whether POC axons grow upon these astroglial cell processes, we performed live imaging using a double transgenic reporter line that distinguished astroglia processes from their associated pathfinding axons *tg(gfap:mCherry-CAAX; gap43:GFP)*. First, we confirmed the earlier timing of pioneering commissural axons crossing the midline of the diencephalon to form the POC (Sup. Mov. 3), which are then followed by axons of the AC crossing the telencephalic midline (Sup. Mov. 4). Using our double transgenic, we conducted a similar timelapse recording from 24 hpf to 28 hpf, which revealed AC axons in direct contact with astroglial cell membranes as they approached the midline from lateral regions (Figure 3, blue boxes) and throughout their track acros the midline to form a commissure (Figure 3, green boxes; Sup Mov 4).

**Figure 3:**
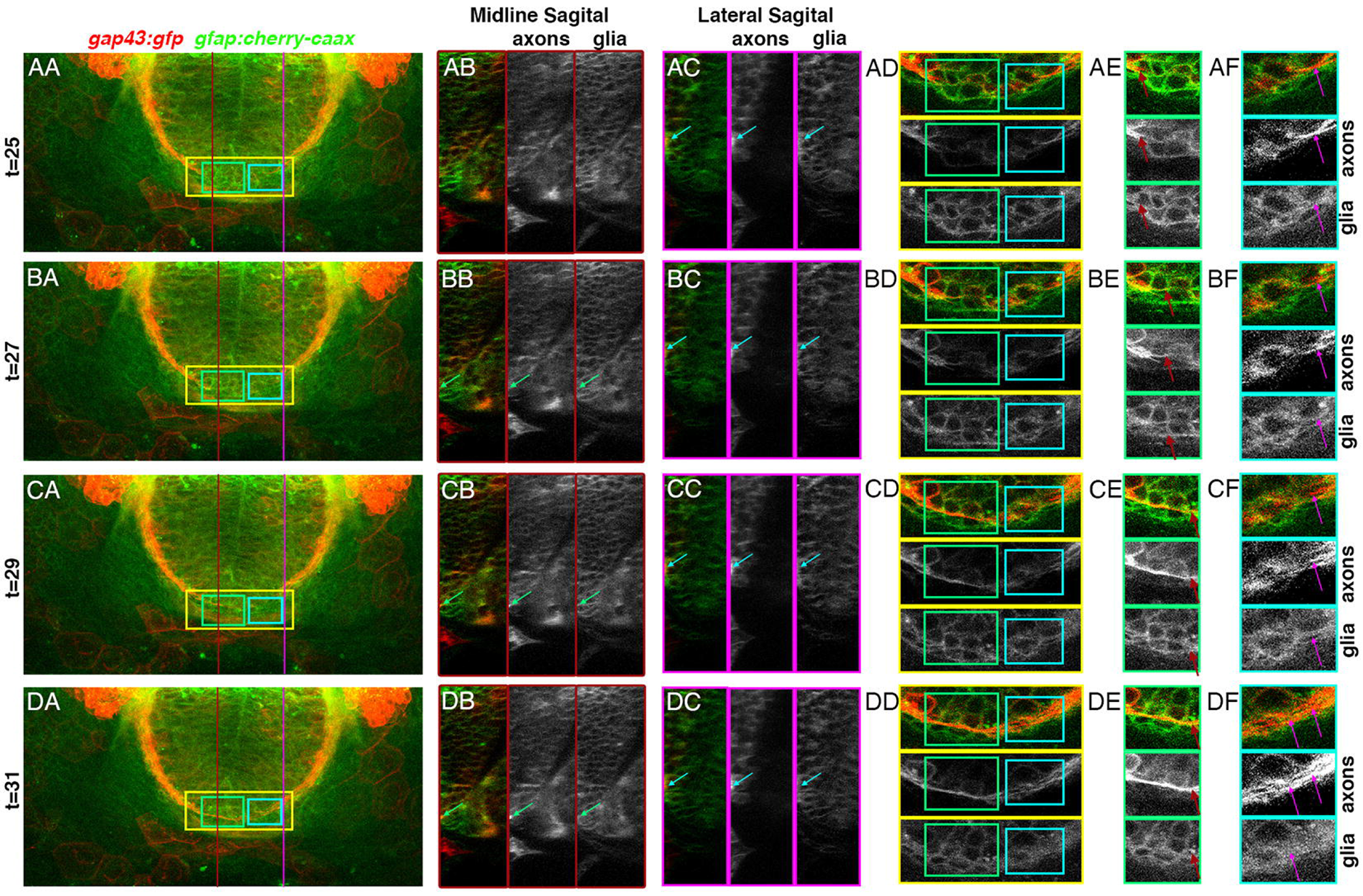
Anterior commissural axons navigate through a meshwork of glial cells while pathfinding towards the midline. AA,BA,CA,DA) MIP color composite dorsal view of the forebrain of a *gfap:cherry-caax* (green); *gap43:gfp* (red) transgenic embryo from 22 hpf to 30 hpf (timestep = 5 minutes). AB, BB, CB, DB) Midline and AC, BC,CC,CD) Lateral and sagittal section of (AA, BA, CA, DA; red and pink lines), respectively, with color composite on left, monochrome *gap43:gfp* center, and *gfap:cherry-caax* on right, showing pioneering axons of the anterior commissure (arrows) pathfinding upon glial surfaces. AD,BD,CD,DD) Dorsal and coronal z-sections of the telencephalon from (AA,BA,CA,DA; yellow boxes), respectively, with selected views (green and blue boxes) highlighting axons pathfinding towards the midline. Selected regions (AE,BE,CE,DE; green boxes) and (AF,BF,CF,DF; blue boxes) from (AD,BD,CD,DD) with color composite *gap43:gfp* (red), and *gfap:cherry-caax* (green) cells (top), monochrome *gap43:gfp* (center), and monochrome *gfap:cherry-caax* (bottom). hpf, hours post fertilization

To date, the characterization of mammalian astrocytic markers, such as antibodies to S100β, Blbp, or GLAST have not proven successful to identify astrocyte subtypes in the zebrafish forebrain. In a previous unbiased screen, four antibodies were found to label clear RGC morphologies in the hindbrain of zebrafish and as such named Zebrafish radial fiber 1 - 4 (Trevarrow, Marks, and Kimmel, 1990). Although we and others have verified that Zrf1 specifically labels zebrafish Gfap (see Figure 4 A,E,I,M; Figure 5 A; (Marcus and Stephen S Easter, 1995)), the antigens recognized by Zrf2-4 remain unidentified. In an effort to determine whether Gfap+ cells in the forebrain represent a heterogeneous population of astroglia, we performed a timecourse of Zrf1-4 expression in the developing zebrafish embryo from 15 hpf to 28 hpf. At all timepoints, anti-Zrf1 labeling completely colocalized with anti-Gfap labeling further verifying the same antigenicity (Figure 4 A,E,I,M). At 15 hpf, prior to any axonogenesis, Zrf1 (Gfap) was consistently found at the more basal extents of the brain, whereas Zrf2-4 were preferentially expressed at the ventricular zone (Figure 4 A-D). By 20 hpf all Zrfs (1-4) were detected in largely overlapping regions of the anterior forebrain (Figure 4 E-H, brackets). Interestingly, at the moment pioneer POC axons start crossing the midline at 23.5 hpf, the positioning of Zrf2-4 in the diencephalon appeared to shift dorsally relative to Zrf1 labeling (Figure 4 I-P, brackets). In contrast, labeling of Zrf2-4 in the telecephalon was much less dense and did not appear to fluctuate over time relative to Zrf1/Gfap. To further investigate the differential positioning of Zrfs in the diencephalon, we conducted higher resolution imaging, which revealed that a greater amount of Zrf2 and Zrf3 preferentially overlapped in the more dorsal regions of the POC, while Zrf4 seemed more tightly concentrated around the POC, and Zrf1/Gfap occupied more ventral portions of the POC region (Figure 5). Additionally, both Zrf2 and Zrf3 appeared to gradually build into a subtle midline peak in the dorsal direction, which correlated with the position of the first optic nerves to cross the midline and form the optic chiasm (Figure 5 E,F, arrowheads).

**Figure 4:**
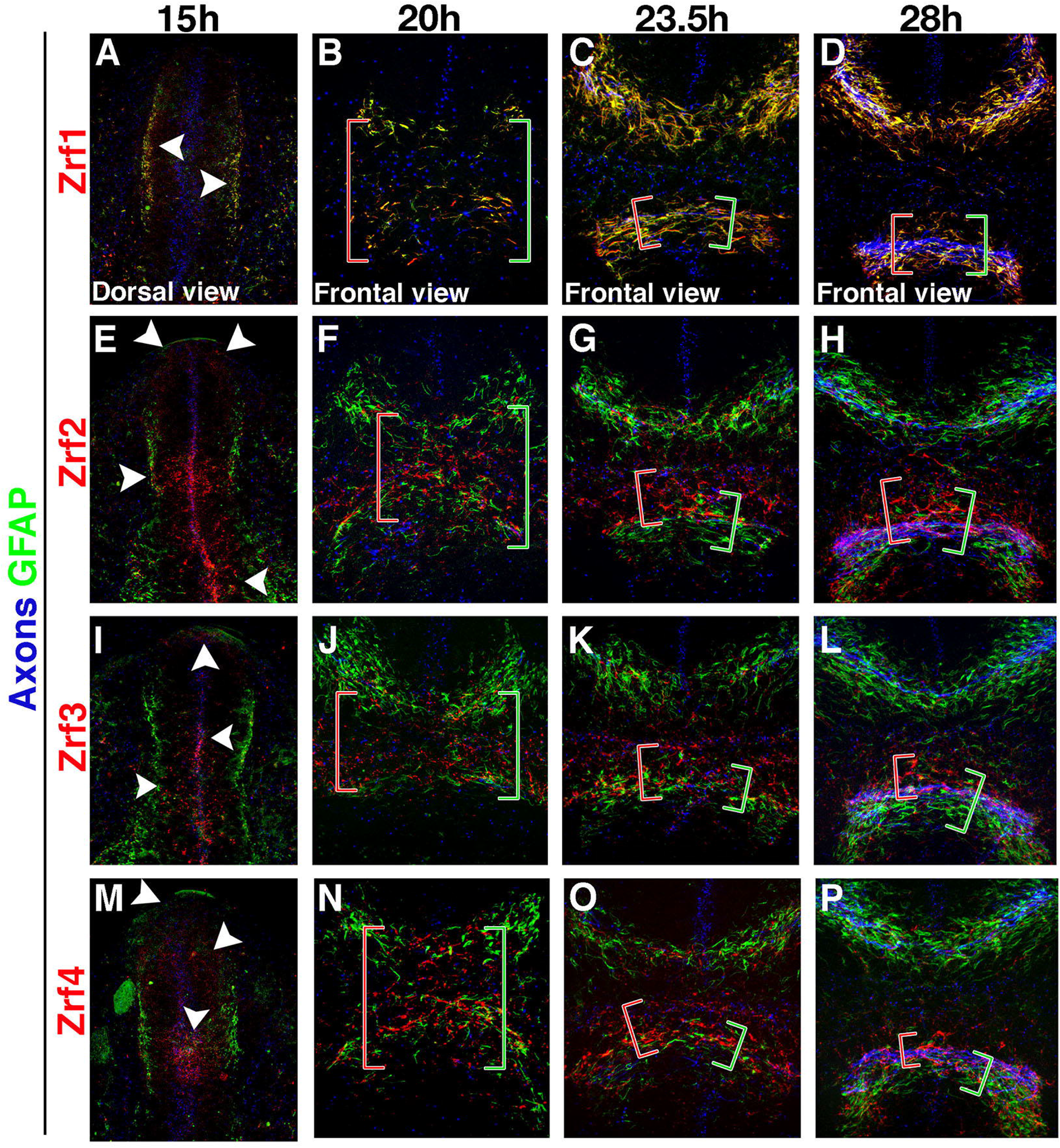
Zebrafish radial fiber markers are differentially labeled across the forebrain. A,E,I,M) Dorsal prosencephalic view of 15 hpf embryos with immunolabeled axons (anti-AT; blue), anti-Gfap (green), and Zrf1-4 (top to bottom; red). B-D, F-G, J-K, N-P) frontal views at 20, 23.5, and 28 hpf, by column. Red brackets: Extent (forebrain B,F,J,N or diencephalon C,D,G,H,K,L,O,P) of red zrf1-4 labeling observed at timepoint. Green brackets: Extent (forebrain B,F,J,N or diencephalon C,D,G,H,K,L,O,P) of Gfap labeling observed at timepoint. hpf, hours post fertilization

**Figure 5:**
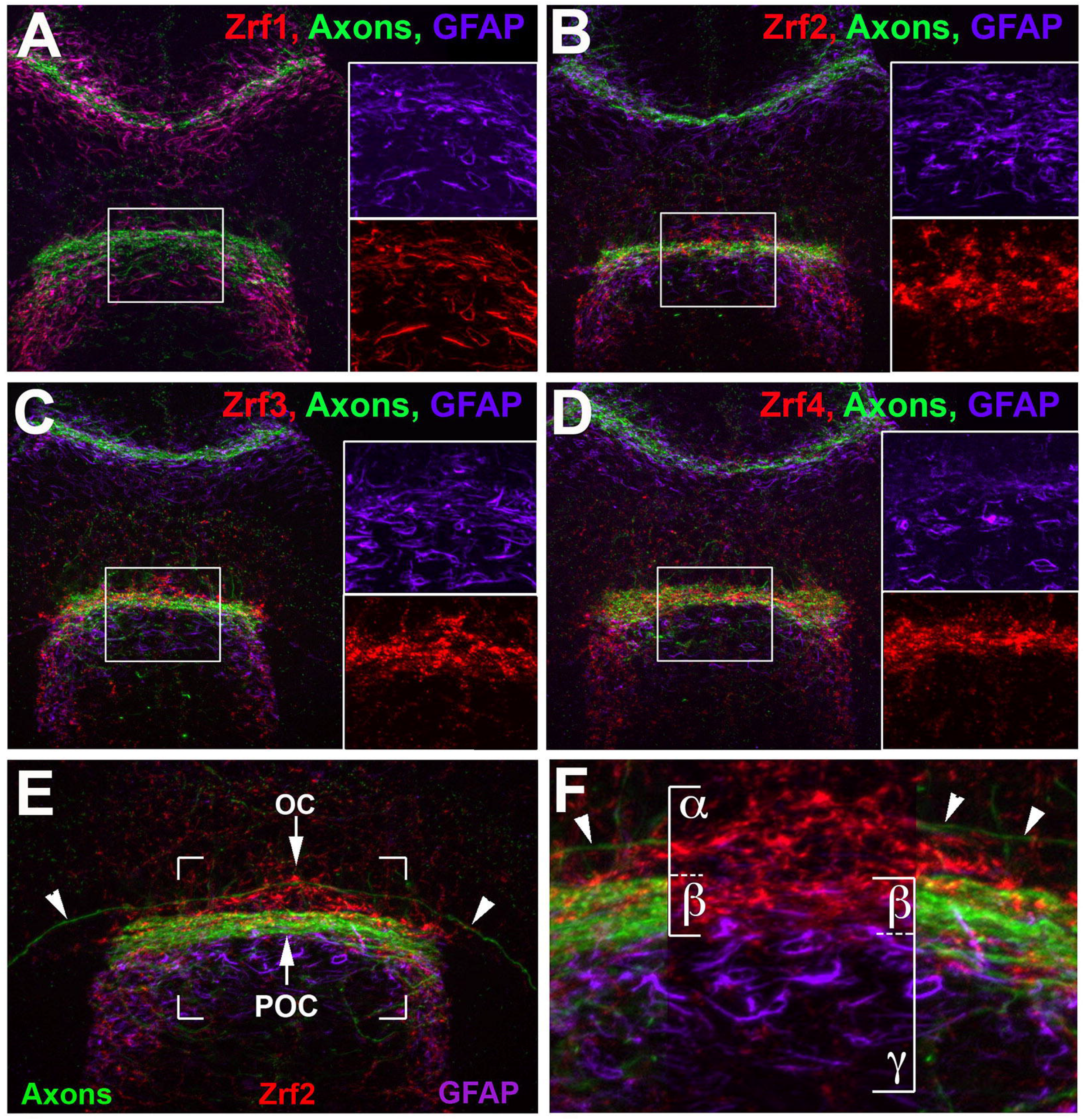
Zrf2-4 labeling is more closely associated with the post optic commissure than Gfap/Zrf1. A-D) Color composite frontal view MIP of 28 hpf embryo labeled with anti-Gfap (purple), anti-AT (green), anti-Zrf1-4 (A-D respectively, red). E) Frontal MIP of 30 hpf embryo with labeled with anti-Gfap (magenta), anti-AT (green), anti-Zrf2 (red), with a selection highlighted in (brackets, F). Note that AT signal at midline region of (F, between brackets) is removed as not to obscure Gfap and Zrf2 labeling. Pioneering OC axons are marked with white arrowheads in (E, F). F) Zrf2 and Gfap overlap in close proximity to the POC (*β*) with Zrf2 labeling extending more dorsally (*α* in comparison to Gfap *γ*). Abbreviations: POC, post optic commissure (white arrow top in E); OC, optic chiasm (White arrow bottom in E). hpf, hours post fertilization

We next used ΔSCOPE to quantitatively assess whether the labeling patterns of Gfap and Zrf2-4 were significantly different from one another. ΔSCOPE is a structural analysis package built for analyzing the 3D structure and composition of the zebrafish POC (Schwartz et al., 2020). In brief, we collected 14 embryos for each of the Zrf markers 2-4, pre-processed them according to the documentation (Schwartz et al., 2020), and compared each Zrf2-4 to either Zrf1/Gfap or to Zrf2. This quantitative analysis confirmed our observations; there was significantly more Zrf2 and Zrf3 signal occupying more dorsal quadrants of the POC as compared to Zrf1/Gfap (Figure 6 A-H). Interestingly, ΔSCOPE did detect some ventral quadrants of the POC to also possess greater amounts of Zrf2 and Zrf3 than that of Zrf1/Gfap (Figure 6 B,F). Although the amount of Zrf4 was quantitatively greater than Zrf1/Gfap at the POC midline, this increase was largely balanced between the dorsal and ventral quadrants (Figure 6 J). Lastly, similar to our qualitative observations, Zrf2-4 showed a statistically-significant tighter association with the POC than Zrf1/Gfap (Figure 7 A-L). There was no difference in the positioning of Zrf3 and Zrf4 around the POC relative to Zrf2. These data, taken together, suggest that Zrf1-4 may each recognize different types of astroglial cells, or label different astroglial subcellular processes that preferentially associate with dorsal or with ventral portions of the POC.

**Figure 6:**
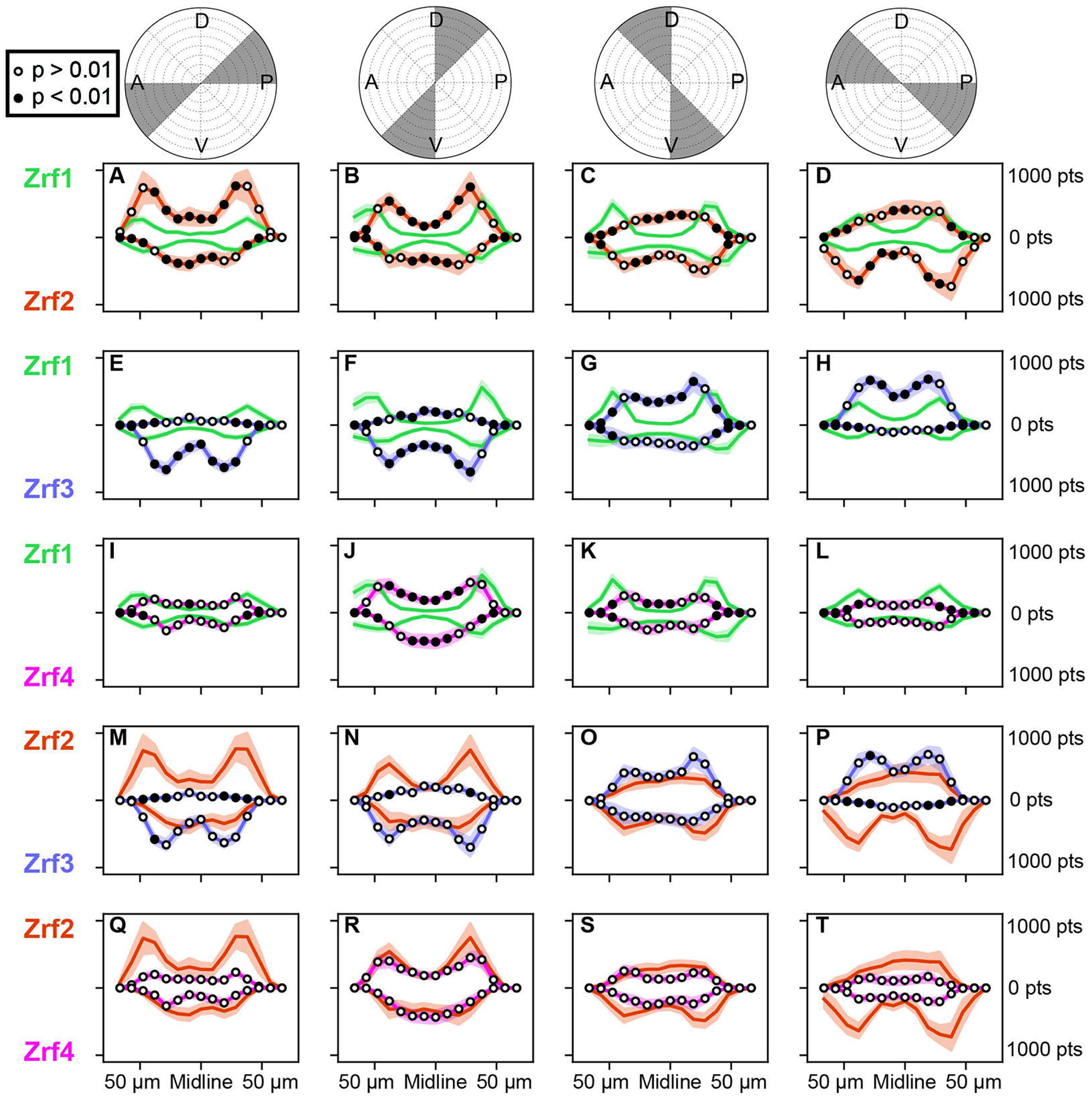
ΔSCOPE analysis of Zrf1-4 expression in the post optic commissure showed differential signal between Gfap and Zrf2-4. A-D, E-H, I-L) Signal comparison between Gfap (n=37) (green) and Zrf2 (n=14) (orange), Zrf3 (n=15) (purple), and Zrf4 (n=12) (pink) respectively showed significantly decreased signal around the commissure relative to Gfap in all quadrants around the POC. This effect was most strongly noted in Zrf2 and Zrf4, where more signal relative to Gfap was observed in the lateral anterior quadrants. M-P, Q-T) Signal comparison between Zrf2 (n=14) (orange) and either Zrf3 (n=15) (purple) or Zrf4 (n=12) (pink) showed no significant differences. hpf, hours post fertilization

**Figure 7:**
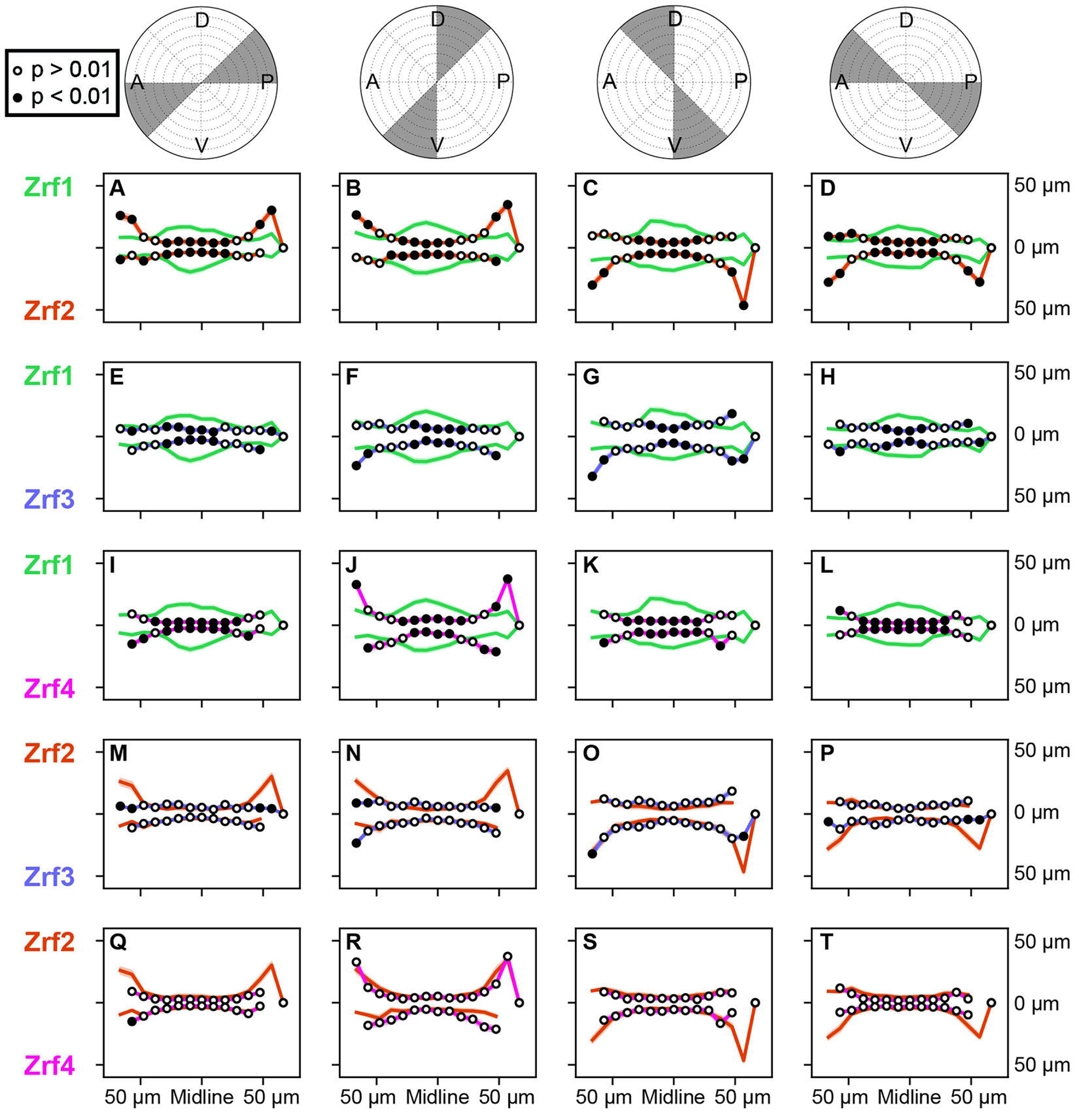
ΔSCOPE analysis of Zrf1-4 expression in the post optic commissure showed greater dorsal localization of Zrf2-4 relative to Gfap. A-D, E-H, I-L) Signal radius comparison between Gfap (n=37) (green) and Zrf2 (n=14) (orange), Zrf3 (n=15) (purple), and Zrf4 (n=12) (pink) respectively showed greater dorsal and posterior localization of relative to Gfap. This was most pronounced in Zrf2 (A-D) but less apparent in Zrf4. M-P, Q-T) Signal comparison between Zrf2 (n=14) (orange) and either Zrf3 (n=15) (purple) or Zrf4 (n=12) (pink) showed no significant differences between Zrf2 and Zrf4 (Q-T) but Zrf3 was noted to have reduced posterior expression. hpf, hours post fertilization

#### 3.3.1 Distinct Glial Morphologies Occupy Different Locations Around The Developing POC

Heterogeneous cellular populations are traditionally first characterized by the identification of distinct cellular subtypes through the presence of different morphologies. Despite the lack of typical mammalian astrocytic markers, unique glial morphologies were discovered in association with the adult zebrafish olfactory bulb that did not conform to the normal radial morphology of radial glia (Scheib and Byrd-Jacobs, 2020; Chapouton et al., 2006). To ascertain whether different astroglial cell morphologies exist in and and are associated with the forebrain commissures in the embryo, we performed gastrula-staged transplantation of cells from the glial reporting lines *tg(gfap:GFP-CAAX; gfap:nls-mCherry)* or *tg(olig2:EGFP)* into wild type embryos and using the gastrula fate map to target cells into the host forebrain (Woo, Shih, and Fraser, 1995). We analyzed the resulting transplants at 28 hpf by immunocytochemistry to label POC axons (Deschene and M. J. Barresi, 2009). We successfully obtained 73 host embryos with isolated colonies of glial cells associating with the developing POC (Figure 8; Figure 9). We took advantage of the *tg(olig2:EGFP)* reporter line to identify and morphologically characterize OPCs residing in the diencephalon. When *olig2:EGFP+* cells were transplanted into gastrula locations targeting the presumptive diencephalon, we were able to visualize a series of cells seated directly dorsal to and intimately contacting the POC (Figure 8 A). We further noted that many of the *olig2:EGFP+cells* associated with the POC exhibited short multi-process contacts with commissural axons (Figure 8 B). These results demonstrate that a subpopulation of OPCs are positioned just dorsal to and interact with post-optic commissural axons, which suggests they may play a supportive guidance role.

**Figure 8:**
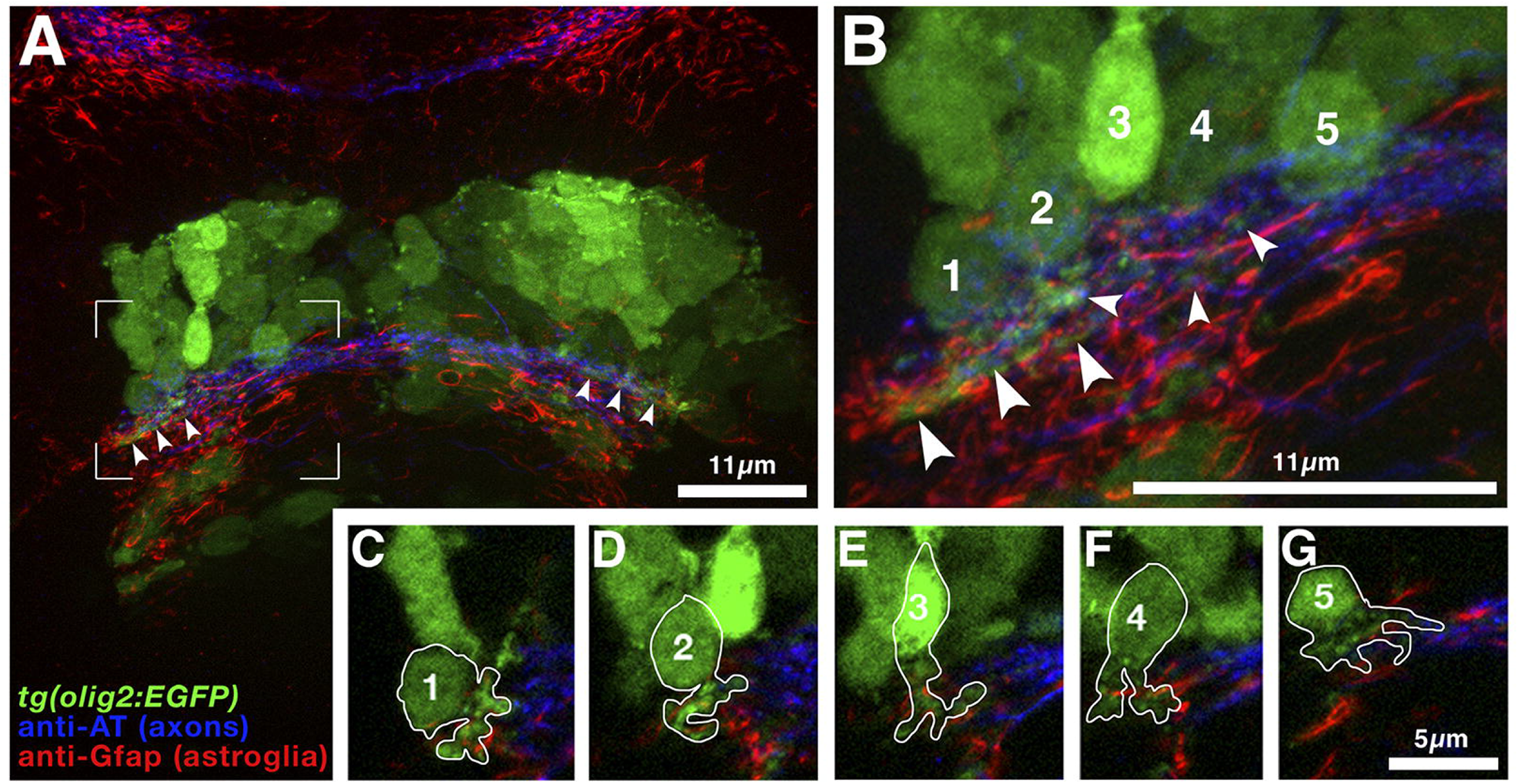
Gastrula stage transplants of *olig2:EGFP+* cells into wild-type zebrafish forebrains revealed the presence of multiple glial morphologies. A) Frontal MIP color composite of zebrafish forebrain at 28 hpf visualizing transplanted *olig2:EGFP+* cells (green) and immunolabeled with anti-Acetylated Tubulin (blue) and anti-Gfap (red); scale bar = 11*μ*m. B) Selection from (brackets, A); scale bar = 11*μ*m. Arrowheads throughout mark *olig2+* processes. C-G) selections of numbered cells from (B) with cell tracing showing varied morphologies; scale bar = 5*μ*m. hpf, hours post fertilization

**Figure 9:**
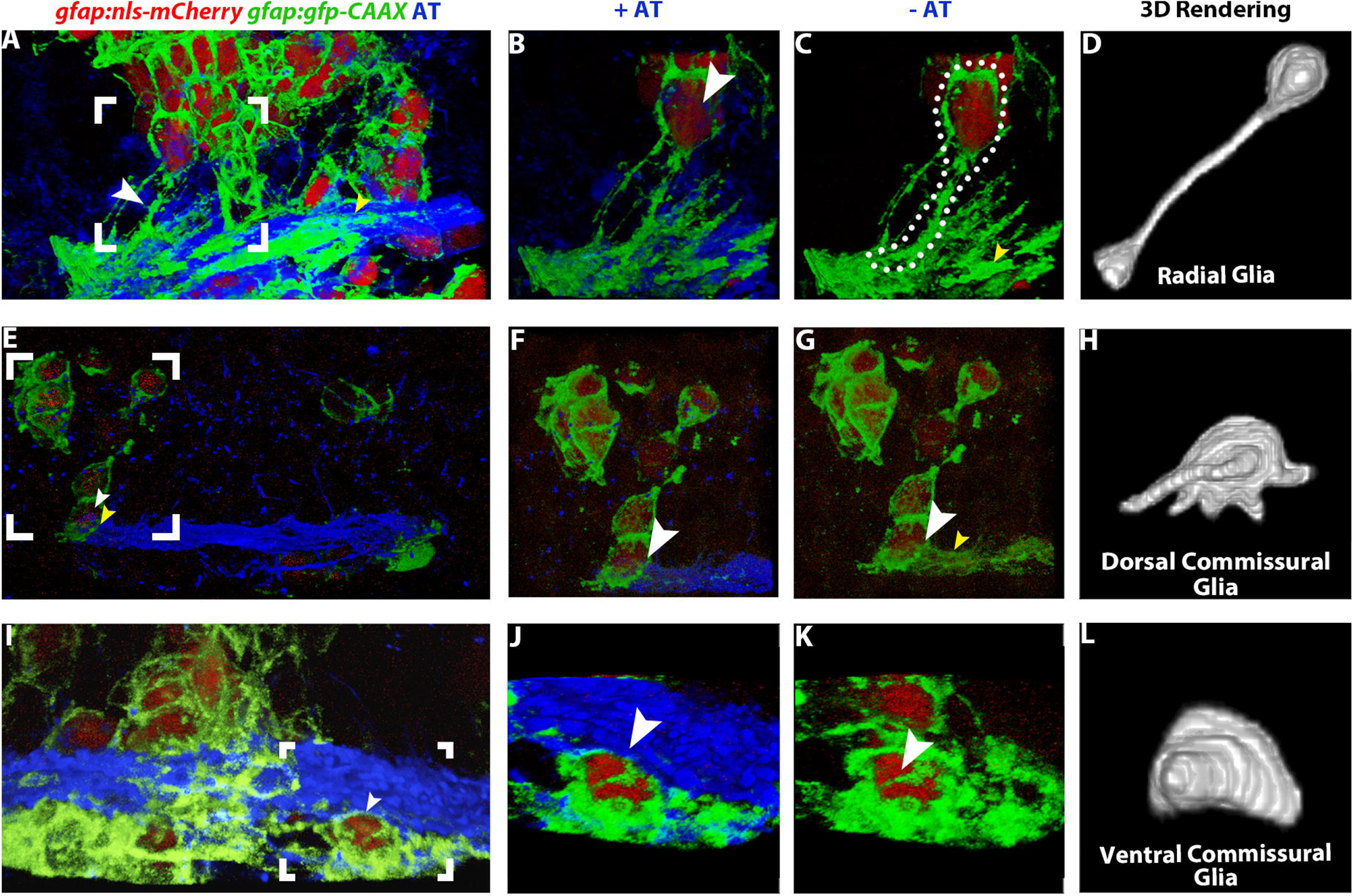
Gastrula stage transplants of *gfap:nls-mcherry;gfap:gfp-caax+* cells into wildtype zebrafish forebrains revealed the presence of multiple glial morphologies. A,E,I) 3D opacity color composite of a 28 hpf forebrain after gastrula stage transplantation of *gfap:nls-mCherry;gfap:gfp-caax+* cells into the prospective forebrain labeled with anti-GFP (green), anti-mCherry (red), and antiAcetylated Tubulin (blue). B, F, J) Selected regions from (A,E,I; brackets), and (C, G, K) same regions without AT shown. White arrowheads indicate highlighted cells with unique glial morphologies and yellow arrowheads indicate glial process in contacts with axons of the POC. D, H, L) 3D renderings of select cells in (D,H,L; white arrows) respectively. hpf, hours post fertilization

We next utilized the *tg(gfap:GFP-CAAX; gfap:nls-mCherry)* reporter lines to characterize the morphology of *gfap-derived* glia in the diencephalon. We were able to detect clear radial glial morphology positioned posterior and dorsal to the POC, with extended end feet that interact with POC axons (Figure 9 C,D, white arrowhead, Sup. Mov. 6 A,D). Moreover, *gfap:nls-mCherry+* progenitor cells can be seen in contact with the radial fiber processes of these radial glial cells (Figure 9). These *gfap+* progenitor cells were also found in direct contact with the dorsal portion of the POC, and showed extended membrane processes that tracked along POC axons (Figure 9 C,E yellow arrowheads, Sup. Mov. 6 B,E). Transplantation of *gfap:nls-mCherry+* cells also showed dense clusters of astroglia along the ventral side of the POC region. These cells had mesenchymal cell morphologies, some of which were in direct contact with POC axons (Figure 9 G, H, white arrowhead, Sup. Mov. 6 C,F). These results suggest that three *gfap+* astroglial morphologies are present surrounding the POC: a radial glial morphology with POC interacting endfeet, a dorsally interacting commissural astroglial cell, and a ventrally interacting population of mesenchymal-like astroglia. To further investigate forebrain glial morphologies, we employed a complementary genetic approach to visually isolate *gfap+* astroglia in the forebrain. By crossing the *tg(ubi:zebrabowM)* transgenic line with the *tg(gfap:CRE^ERT2^*) estrogen-responsive CRE transgenic line, we were able to clonally label *gfap+* cells following Tamoxifen treatment to induce recombination of zebrabow’s fluorescent cassette (Pan et al., 2005; Jungke et al., 2015). This approach produced fluorescently distinct cell bodies seated adjacent to the POC, similarly positioned to those observed following our *gfap:nls-mCherry* gastrula cell transplantations (Figure 10 A,B). Although the cytoplasmic accumulation of the zebrabow GFP fluorophore expression was insufficient to completely fill cell processes, we were able to define their intimate proximity with Gfap protein localization, which corroborated the glial identity of these clones (Figure 10 C,D). During this analysis, we also noted close association between *gfap+* clones and AT+ axons (Figure 10 C-F). Additionally, when we employed 3D rendering analysis of these recombined *gfap+* clones, we were able to visualize clear radial glial morphology spanning from the midline ventricle to pial surfaces in the commissure regions (Figure 10 G). Moreover, immunolabeling of 28 hpf embryos with Tamoxifen-induced gfap+ clones for AT and Gfap protein expression demonstrated direct interactions between POC axons with astroglia cells (Figure 10 H-J). These results confirm *gfap+* astroglial cells provide a direct substrate for pathfinding commissural axons in the forebrain.

**Figure 10:**
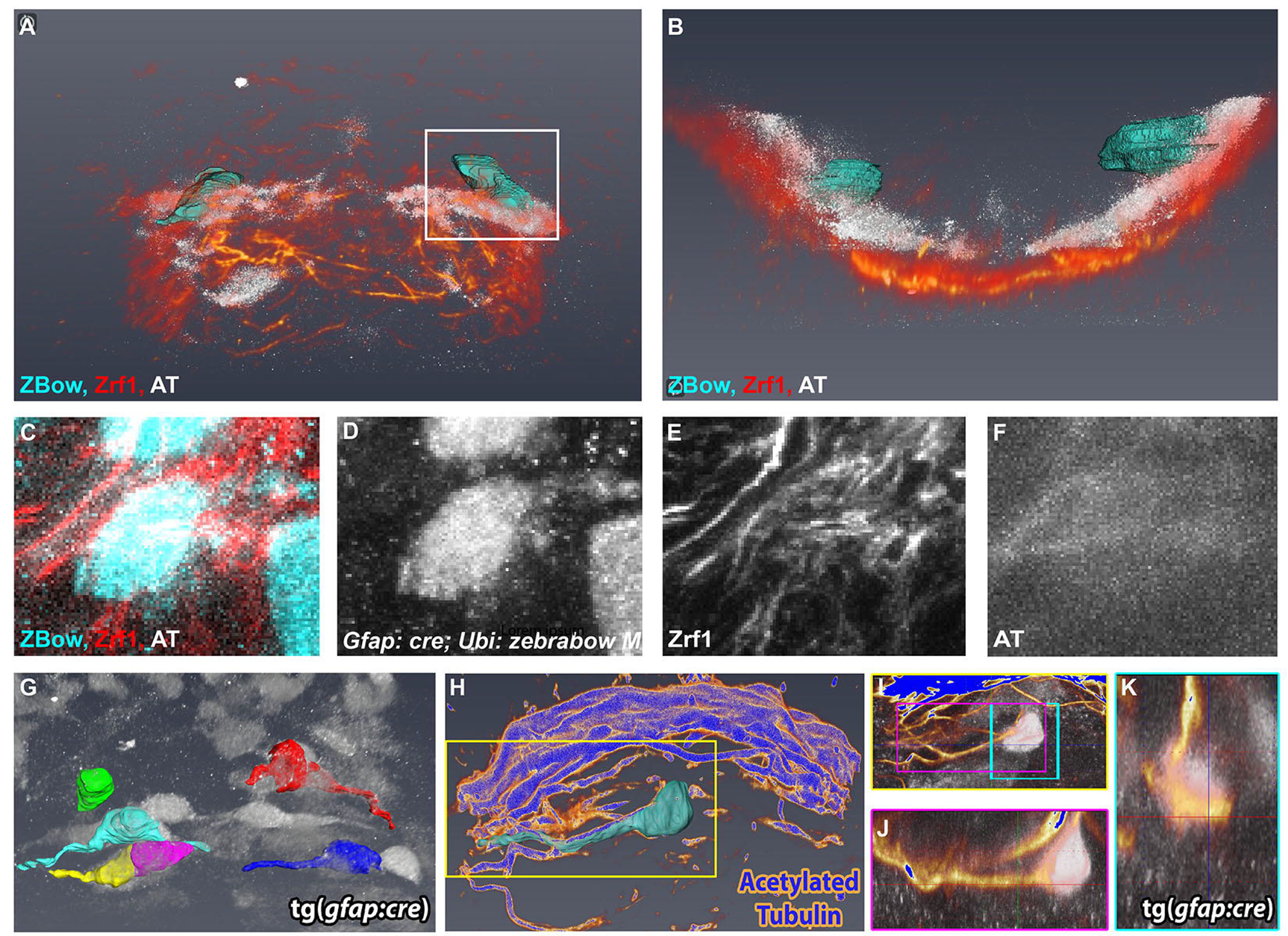
Clonal analysis of glial cells show pathfinding post optic commissural axons migrate medially along glial processes. A,B) 3D opacity color composite of tamoxiphen treated *tg(ubi:zebrabow-M;gfap:cre^ERT2^*) 28hpf embryo labeled with anti-GFP (cyan), anti-acetylated tubulin (white), and anti-Gfap (Zrf1, red), showing frontal (A), and anterior (B) views of the POC region. C-F) MIP color composite (C) and single channel (*ubi:zebrabow-M;gfap:cre*ERT2** (D), Zrf1 (E), acetylated-tubulin (F)) views of (A, inset). G) 3D opacity of 28 hpf tamoxiphen treated *tg(ubi:zebrabow-M;gfap:cre^ERT2^)* embryo immunolabeled for anti-Gfp (white). G) 3D rendered *gfap+* glial cells exhibited radial like morphology along the embryo midline. H-J) 3D opacity of tamoxiphen treated *tg(ubi:zebrabow-M;gfap:cre^ERT2^)* embryo labeled with anti-Acetylated Tubulin (blue-gold) and 3D rendered glial cell from anti-Gfp labeling (cyan). I) Frontal inset from (H). J) Magenta inset from (I). K) Sagital MIP inset from (I, cyan). hpf, hours post fertilization

#### 3.3.2 Oligodendroglial Progenitors Interact with Commissural Axons

Olig2 progenitor cells (OPCs) play an important role in the generation of both neural and glial cell types in the developing CNS (Fontenas and Kucenas, 2018; Kriegstein and Alvarez-Buylla, 2009a; Borodovsky et al., 2009). Although the ontogeny of OPCs in the spinal cord has been extensively studied (Mitew et al., 2014; Ravanelli and Appel, 2015), less is known about the lineage of, and requirements for, OPCs during forebrain development. Importantly, evidence has shown that *olig2+* cells give rise to myelinating cells of the spinal cord by 120 hpf (Pogoda et al., 2006). Based on our 48 hpf characterization of *tg(olig2:EGFP)* expression and related chimeric analyses demonstrating that *olig2+* cells intimately contact commissural axons, we hypothesized that OPCs in the forebrain play essential roles in influencing the successful development of those same forebrain commissures. To investigate this possibility, we sought to characterize the temporal and spatial patterning of *olig2+* cell populations during early commissure formation.

We used the *tg(olig2:EGFP)* transgenic line to visualize OPCs in the forebrain from 20 to 36 hpf, capturing the full embryonic range of commissure development. By 20 hpf, *olig2:EGFP* expression was observed predominantly in the diencephalon and was represented by the bilateral positioning of two qualitatively different cell populations. The most dense *olig2:EGFP+* cellular populations occupied the lateral regions of the diencephalon (Figure 11 A, white brackets, Sup. Mov. 6 A), while a sparse row of cells was positioned more medially, and spanned the midline (Figure 11 A, inset, asterisks). The medially-positioned *olig2:EGFP+* cells exhibited ventrolaterally extending processes along the presumptive pathway of the future POC (Figure 11 A, inset arrowheads). Additionally, a small population of bilaterally symmetrical *olig2:EGFP+* cells were seen in the telencephalon in the position of the presumptive anterior commissure (Figure 11 A, arrows).

**Figure 11:**
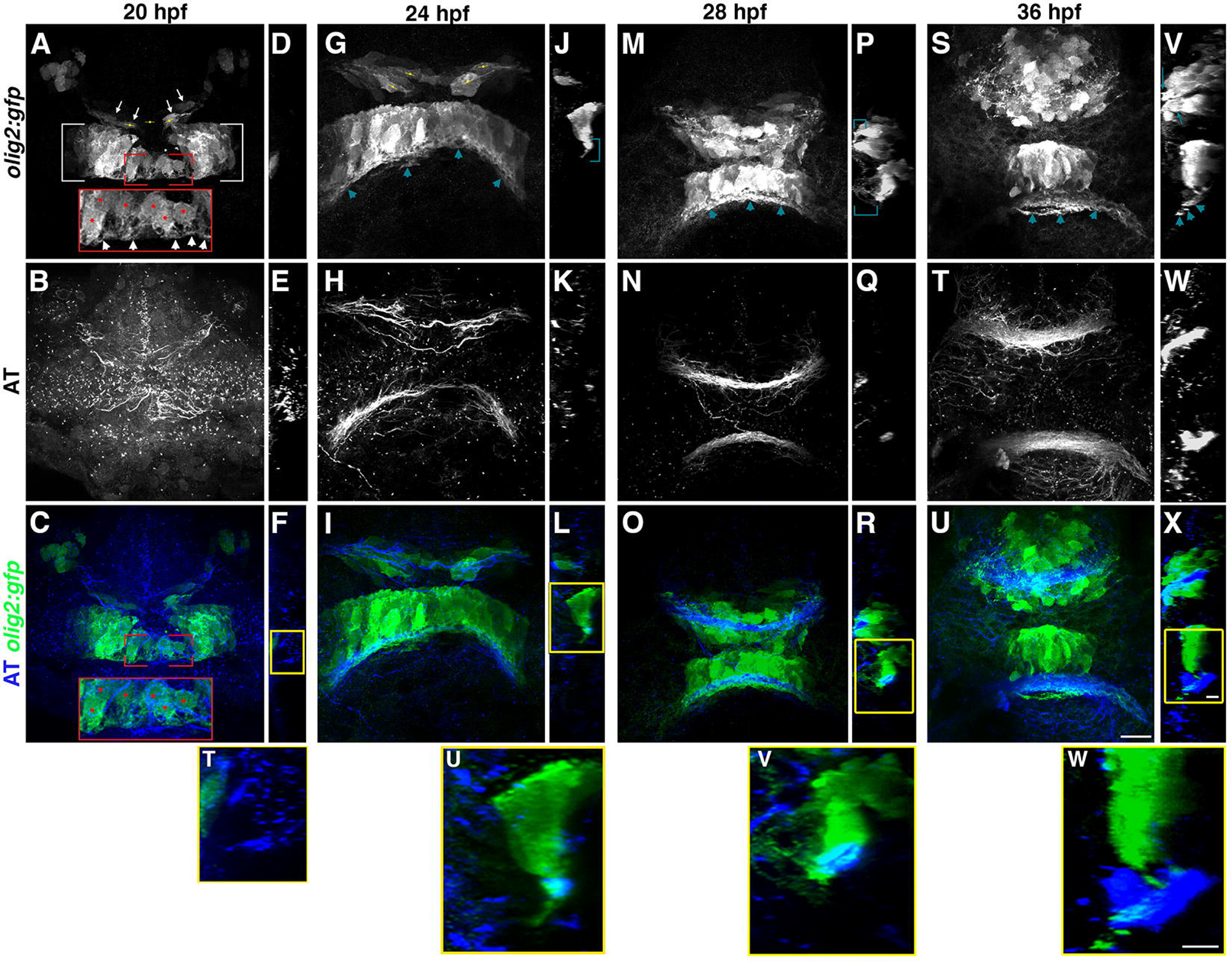
Oligodendrocyte progenitor cells and *olig2+* cells progressively populate the preoptic area and the tract of the anterior commissure where they protrude and make contact with the developing commissures. A-C,G-I,S-U) frontal view of *tg(olig2:EGFP)* (monochrome: A,G,M,S; green: C,I,O,U) embryo labeled with anti-AT (monochrome: B,H,N,T; blue: C,I,O,U), at 20, 24, 28, and 36 hpf, by column (scale bar = 11*μ*m). D-F,P-R,V-X) midline sagittal sections of (A-C,G-I,S-U), respectively (scale bar = 4.7*μ*m). Inset in (A,C) from red bracketed regions, with red asterisks indicating *olig2+*cells adjacent to POC and (A white arrowheads; G,M,S,V teal arrowheads) mark *olig2+* processes extending into POC tract (V teal arrows) and AC. T-W) Yellow boxed insets from (F,L,R,X; scale bar = 4.7*μ*m). Yellow axis markers (A,G,M) indicate observed oligodendrocyte polarity. hpf, hours post fertilization

Between 20 and 36 hpf we noticed an obvious qualitative increase in *olig2:EGFP+* cells in both the telencephalon and diencephalon (Figure 11). Most notable was the clear increase in the density of *olig2:EGFP+* cells within the telencephalon over time, which became concentrated fully around the anterior commissure. Although we also noticed an increase in the number of *olig2:EGFP+* cells in the medial portion of the diencephalon, the overall expressing population became positionally more focused upon the medial region by 36hpf. As suggested by our 48 hpf characterization of *olig2:EGFP+* cells, this coalescing of *olig2:EGFP+* cells at the midline just above the POC appears to continue through day two of commissure development (See Figure 1 V-Z). Furthermore, the *olig2:EGFP+*cell processes evident at 20 hpf remained a consistent structure throughout the time course. Moreover, these processes increased in thickness and number while remaining largely associated with the developing anterior and post optic commissures (Figure 11 G, M, S, teal annotations).

In addition to the extension of membranous processes to the POC and AC, we also observed evidence of changes to *olig2:EGFP+cell* morphology in the telencephalon as well as to their peripheral processes over time. Between 20 and 24 hpf, *olig2:EGFP+* cells showed a bipolar protrusive morphology oriented nearly perpendicular to the midline and parallel to the future anterior commissure axis (Figure 11 A,G, telencephalon, yellow axis markers, Sup. Mov. 6 B,C). However, by 28 hpf the orientation of these telencephalic *olig2:EGFP+*cells became more varied and displayed numerous membrane extensions toward and associated with the anterior commissure, a phenotype which had intensified by 36 hpf (Figure 11 M,P,S,V, teal annotations, Sup. Mov. 6 D). Importantly, the *olig2:EGFP+* cell projections in both the diencephalon and telencephalon were predominantly intermixed with the developing tracts and commissures for both the POC and AC throughout the timecourse (Figure 11 C,I,O,U, Sup. Mov. 6 A-D). These results demonstrate a progressive elaboration of *olig2:EGFP+* cells are positioned in both the diencephalon and telencephalon and their cell processes interact with commissural axons.

#### 3.3.3 Function of Radial glia and OPCs as stem cells in the forebrain

To examine whether radial glial cells and OPCs share both a lineage and functional relationship as forebrain stem cells, we characterized cell division in *tg(olig2:EGFP; gfap:nls-mCherry)* double transgenic embryos, using the M-phase marker anti-Phospho-Histone H3 (pH3) to label cells in mitosis. At 22 and 24 hpf, the *olig2:EGFP+* cells in the anterior most midline region of the diencephalon were largely negative for *gfap:nls-mCherry* expression (Figure 12 A,F,I,N, Sup. Mov. 7 A,B). Double positive *olig2:EGFP* and *gfap:nls-mCherry* expressing cells were consistently observed further from the midline in more posterior positions of the forebrain (Figure 12 B,J,R). However, by 28 hpf, *olig2:EGFP* and *gfap:nls-mCherry* double positive cells can be seen at the diencephalic midline (Figure 12 Q,R, Sup. Mov. 7 C). In the telencephalon, a vast majority of the *olig2:GFP+* cells developed later than those detected in the diencephalon over this same time course, and the *olig2:EGFP+*cells were all double positive for *gfap:nls-mCherry* expression (Figure 12 F,G,N,O,V,W).

**Figure 12:**
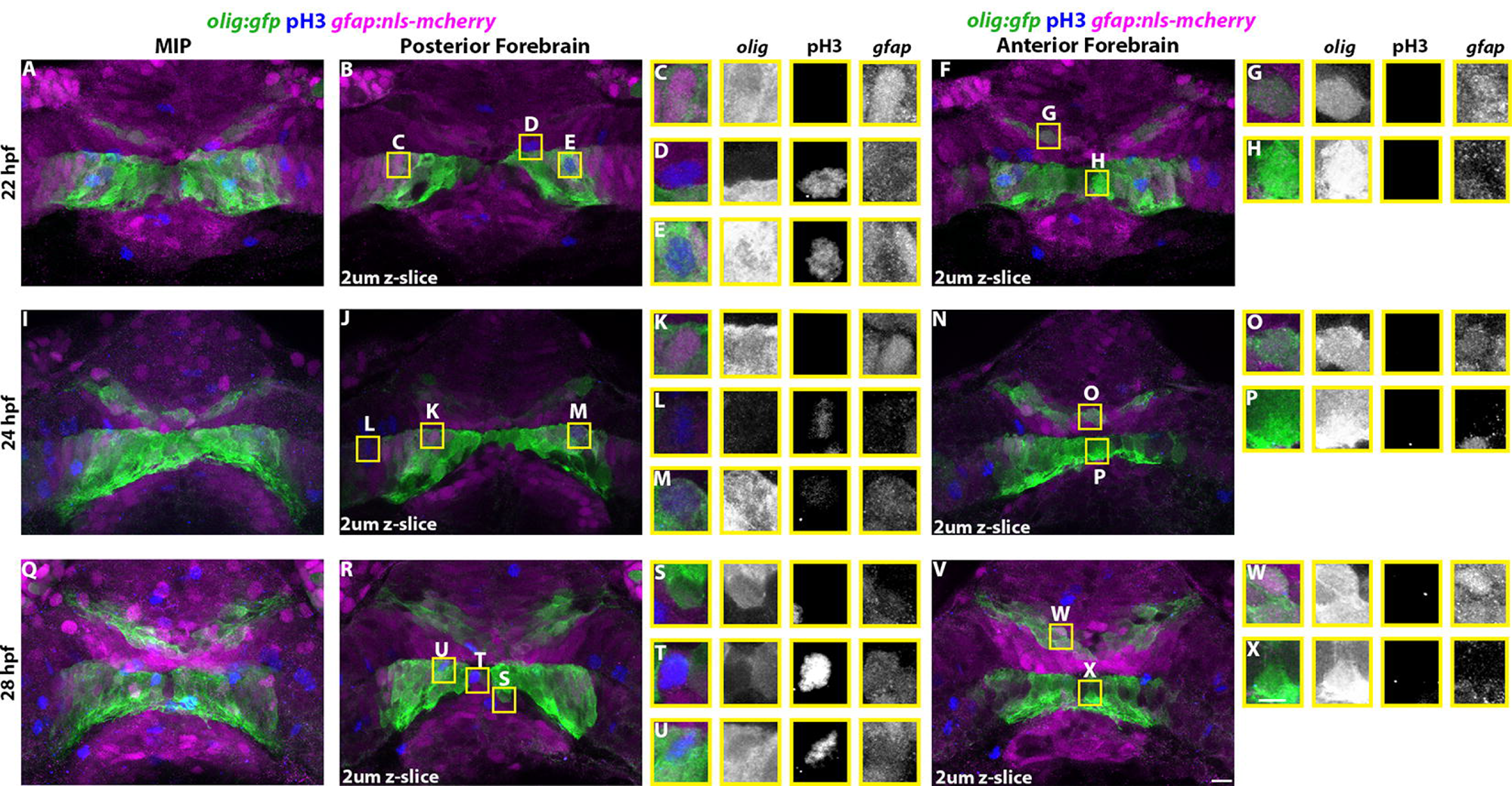
The glial progenitor pools of the developing telencephalon and diencephalon exhibit their own distinct lineage and spatial-temporal restrictions. A,I,Q) Frontal color composite MIP and posterior (B,J,R) and anterior (F,N,V) selected 2*μ*m z slices of *tg(gfap:nls-mcherry* (magenta); *olig2:gfp* (green)) embryos labeled for pH3 (blue) at 22 (A,B,F), 24 (I,J,N), and 28 hpf (Q,R,V). Individual selected cell inset composites from (B, J, R, F, N, V) are shown in (C-E, K-M, S-U, G-H, O-P, W-X) with single channel monochromes of *olig2:gfp*, anti-pH3, and *gfap:nls-mCherry*, from left to right. Scale bar = 11*μ*m. hpf, hours post fertilization

Upon characterization of cell division patterns in the forebrain, we primarily observed that the PH3 label was restricted to the midline and lateral ventricles (Figure 12 D,E,L,M,T,U), which is consistent with current models of nuclear translocation to the apical surface of the ventricles during progenitor cell division (Götz and Barde, 2005; Buckley and Clarke, 2014). Interestingly, radial glia or OPCs that were pH3+ only expressed *gfap:nls-mCherry* or *olig2:EGFP* respectively and were never double positive for the expression of both transgenes (Figure 12). This seems to suggest that PH3+ cells that are *gfap:nls-mCherry+* and *olig2:EGFP* - or *olig2:EGFP+* and *gfap:nls-mCherry*-have different stem cell roles in the forebrain. We hypothesize that *olig2:EGFP* and *gfap:nls-mCherry* double positive cells either divided at a much slower rate, or had otherwise adopted a more differentiated and less proliferative state.

### 3.4 Characterizing Cell Differentiation in the Forebrain

In an effort to elucidate the differentiated cell fates that radial glia and OPCs adopt in the developing forebrain, we characterized the association of neuronal and myelinating glial cell types relative to known progenitor cell types. Our earlier characterization of *olig2:EGFP+*cell morphology indicated a resemblance to oligodendroglial cells with processes that were associated with commissural axons, which is a behavior suggestive of a possible myelinating function. Glial cells that co-express *nkx2.2* have been demonstrated to adopt myelinating oligodendrocyte cell fates (Mitew et al., 2014; Zhou, Choi, and Anderson, 2001). To investigate whether *olig2:EGFP+* cells in the forebrain become myelinating glia, we tested for the dual expression of *nkx2.2:GFP* with *olig2:mCherry* at 28 and 36 hpf. Although the number of cells expressing *nkx2.2:GFP* increased dramatically over this time period, we did not observe any colocalization of *nkx2.2:GFP* in *olig2:EGFP+* cells (Figure 13 A,E, Sup. Mov. A,B). Therefore, despite the oligodendroglial morphology (long axon-interacting processes) displayed by *olig2:EGFP+* cells in the forebrain, lack of *nkx2.2* colocalization suggests oligodendrocyte differentiation has not yet begun. This is consistent with previous reports indicating that the specification of myelinating oligodendrocytes occurs later in larval development (initiate at 2 days post fertilization, myelin positive at 3 dpf (Ackerman and Monk, 2016; Fontenas and Kucenas, 2018; Kucenas, Snell, and Appel, 2008).

**Figure 13:**
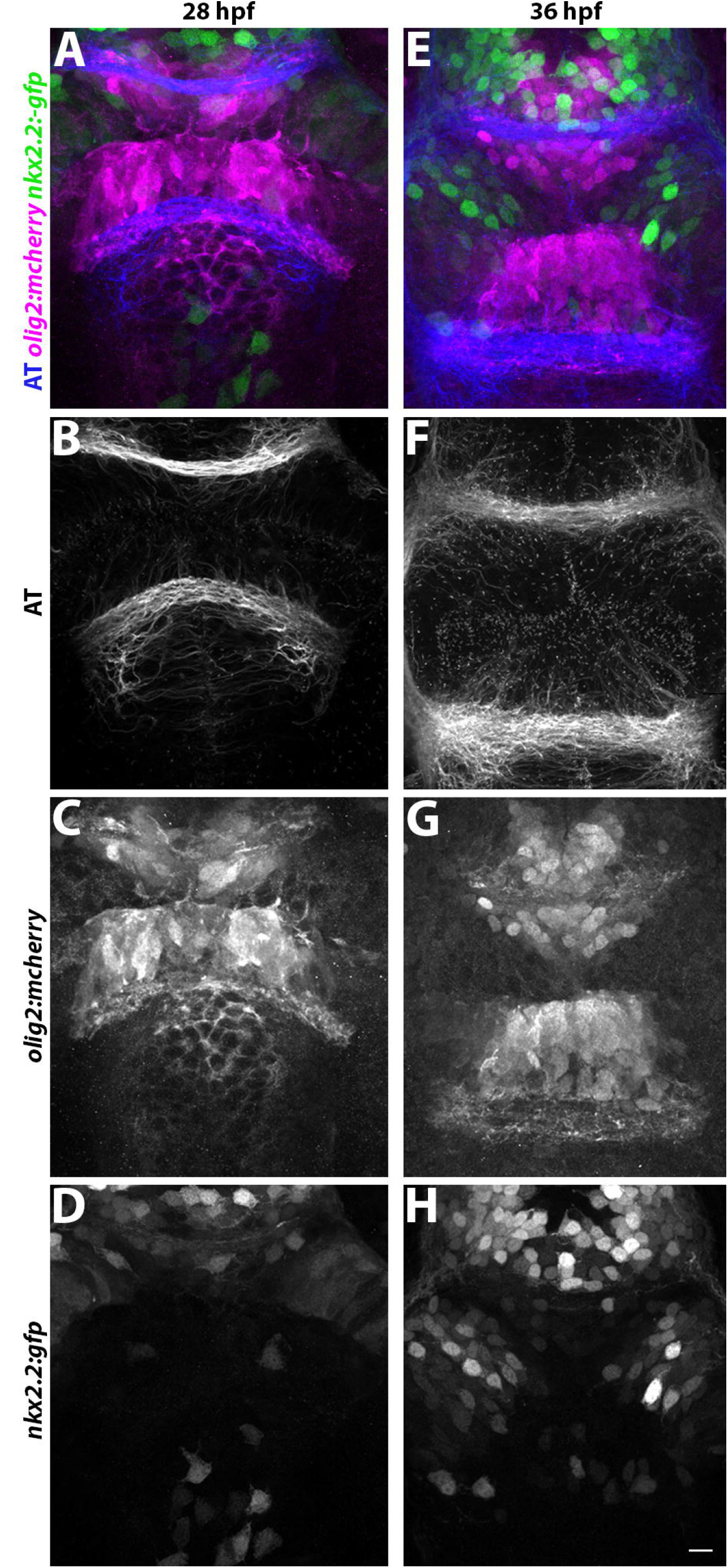
*olig2+* cells do not adopt a myelinating fate by 36 hpf despite widespread expression of *nkx2.2* in surrounding cells. A,E) Frontal color composite MIP of *tg(nxk2.2:gfp* (green); *olig2:mCherry (magenta))* labeled with AT (blue) at 28 and 36 hpf, respectively. Monochrome single channels MIPs of (A,E) by column: (B,F) anti-AT, (C,G) *olig2:mCherry*, (D,H) *nkx2.2:gfp*. Scale bar = 11*μ*m. hpf, hours post fertilization

Both *olig2-* and *gfap*-expressing cells have been shown to give rise to neuronal populations in the spinal cord (Kriegstein and Alvarez-Buylla, 2009a; Johnson et al., 2016; H. Kim et al., 2008; Lyons and Talbot, 2015); therefore we next investigated whether the identified progenitor populations of *olig2:EGFP+* and *gfap:nls:mCherry+* were contributing to neuronal populations in the forebrain during commissure formation. Using the *tg(huC:GFP)* neuronal transgenic reporter line, we observed distinct neuronal populations in both the diencephalon and telencephalon at 24 hpf, which increased in number by 28 hpf and again by 36 hpf (Figure 14). These *huC:GFP+* neuronal populations were initially most dense around the anterior commissure with very little neuronal differentiation present in the diencephalon at 24 hpf (Figure 14 A). Although the number of *huC:GFP+* neurons grew around both commissures, a vast majority of *huC:GFP+* axons contributed directly to the anterior commissure (Figure 14, E, H insets, arrows) as compared to huC:GFP+ contributions to the POC (Figure 14 H, arrowheads).

**Figure 14:**
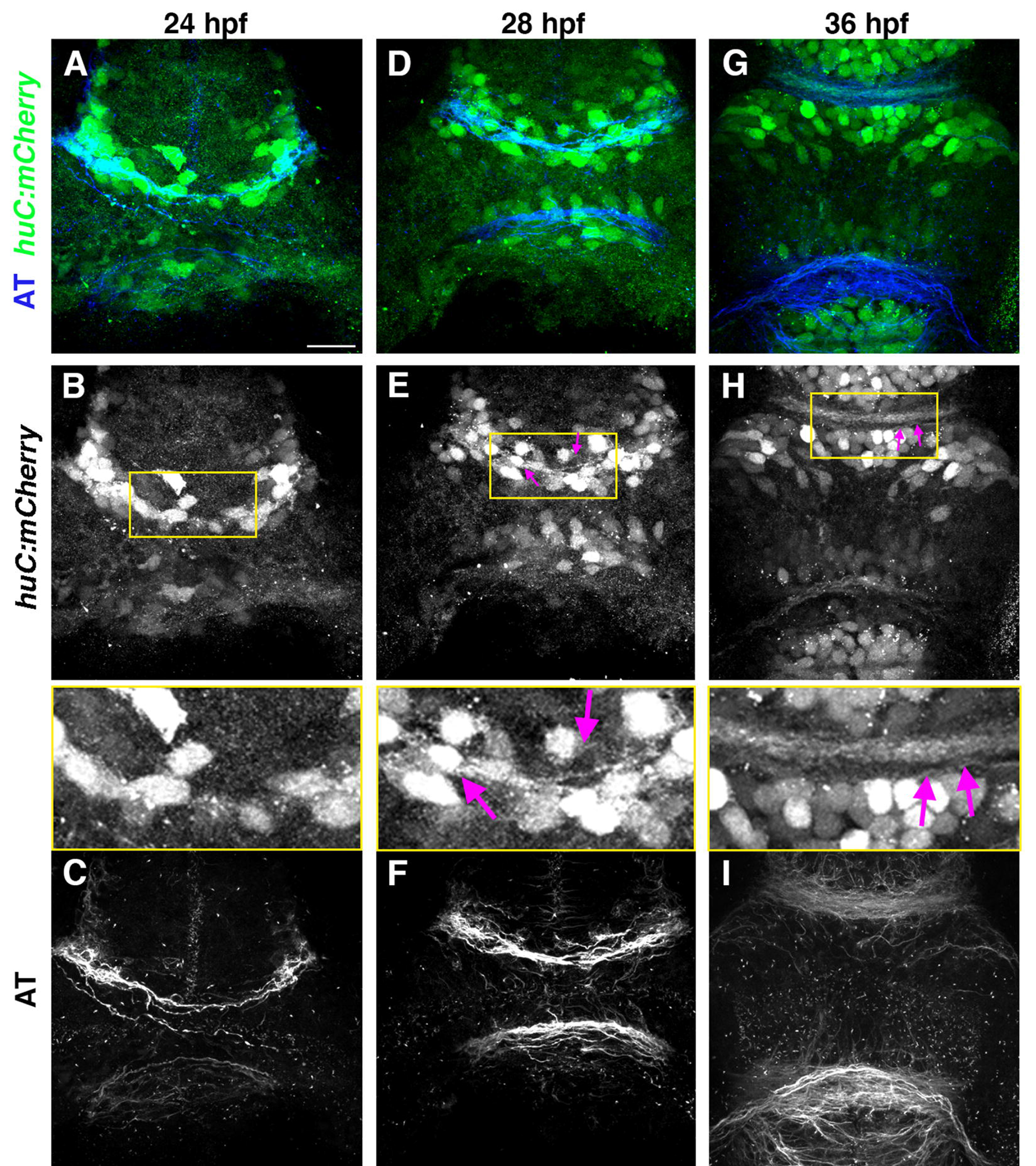
Forebrain *huC+* neurons progressively populate the regions directly adjacent to the anterior and post optic commissure, and progressively contribute to anterior commissure axons. A,D,G) Color composite frontal MIPs of *tg(huC:mCherry)* (green) embryos labeled with anti-AT (blue) at 24, 28, 36 hpf respectively (scale bar = 11*μ*m). Single channel monochromes of (A,D,G) are *huC:mCherry* (B,E,H) and AT (C,F,I). Enlarged yellow boxed insets of (B,E,H) highlight the AC region. Pink arrows indicate *hwC*+processes extending into the AC. hpf, hours post fertilization

To test whether any of these neuronal populations are derived specifically from *gfap+* or *olig2+* stem cell lineages, we performed anti-HuC/D immunolabeling in the double *tg(olig2:EGFP; gfap:nls-mCherry)* backgrounds at 24 and 28 hpf. In both the telencephalon and diencephalon, we discovered examples of all four possible neuronal lineages: HuC/D+ cells devoid of any transgene expression, HuC/D+ cells that were co-expressing both *olig2:EGFP* and *gfap:nls-mCherry*, as well as separate HuC/D+ cells double labeled for expression with either the *olig2:GFP* or *gfap:nls-mCherry* transgenes (Figure 15, Sup. Mov. 9 A,B). Interestingly, there were some clear qualitative distinctions observed over this time course. At 24 hpf there were fewer HuC/D+ neurons in the diencephalon as compared to the telencephalon, yet a majority of these HuC/D+ cells were colabeled with *gfap:nls-mCherry* or devoid of any transgene expression (Figure 15 C). However, by 28 hpf the telencephalic HuC/D+ neurons positioned dorsal to the developing anterior commissure were predominantly *olig2:EGFP+* (Figure 15 I,K), whereas those HuC/D+ neurons positioned more ventral to the anterior commissure continued to primarily co-express *gfap:nls-mCherry+* (Figure 15 I,J).

**Figure 15:**
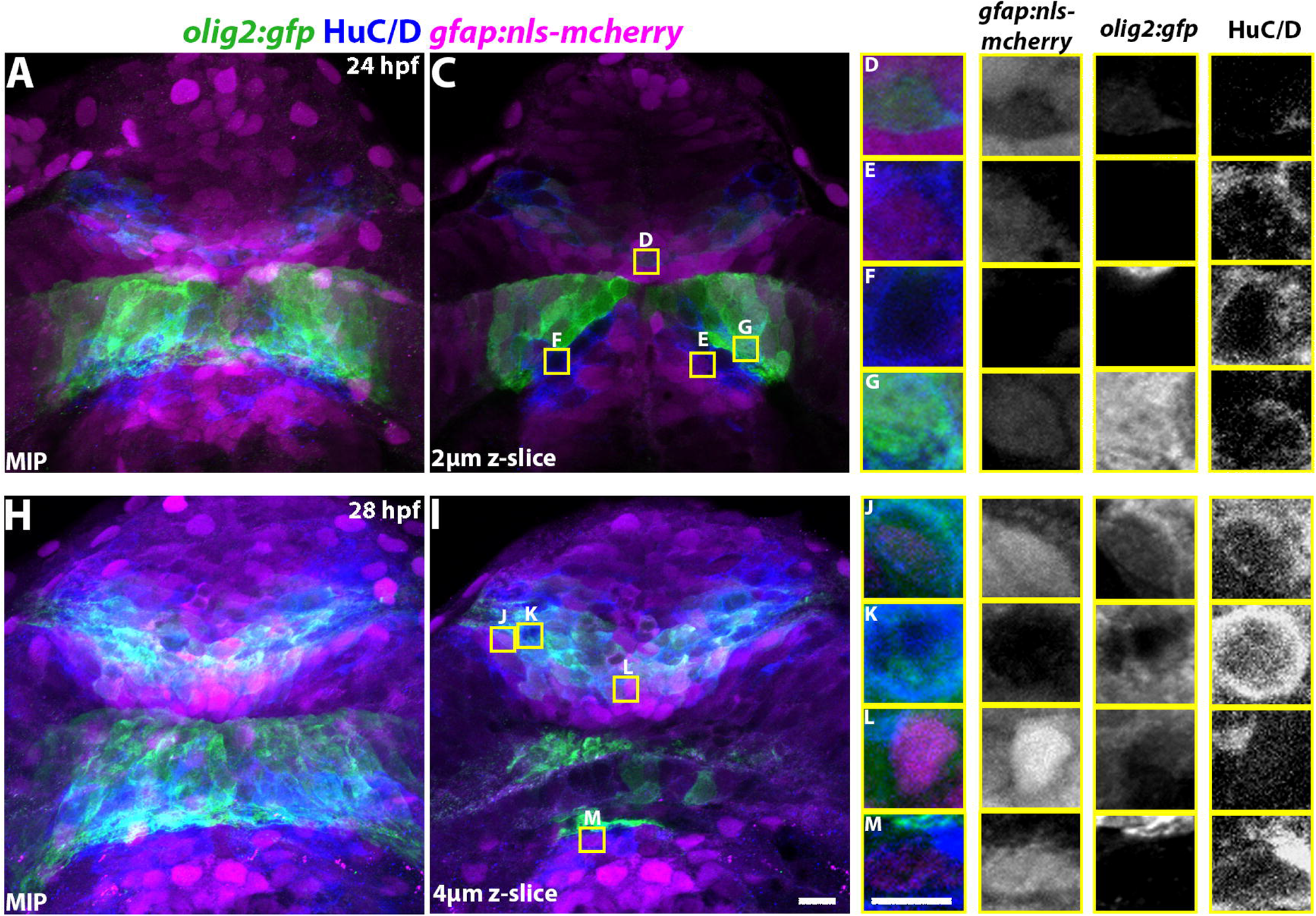
Radial glia and oligodendrocyte progenitor cells exhibit differential neuro-progenitor contributions in the diencephalon and telencephalon. A,H) Color composite frontal MIP with coronal sections (C,I) of *tg(gfap:nls-mcherry)* (magenta); *tg(olig2:EGFP)* (green) embryos labeled with anti-HuC/D (blue) at 24 hpf (A,C) and 28 hpf (H,I). C,I) Individual selected cell inset composites from (D-G, J-M) with single channel monochromes of *gfap:nls-mCherry, olig2:EGFP*, and anti-HuC/D, from left to right. Scale bar = 11*μ*m. hpf, hours post fertilization

To further examine the potential fate of these HuC/D+ neuronal cells in the forebrain, we evaluated these populations for the co-incidence of *nkx2.2* HuC/D expression. While in glial cells *nkx2.2* is a marker for myelinating cell identities, its expression in neurons denotes specification toward serotonergic fates (Panula et al., 2010; Cheng et al., 2003). Interestingly, the majority of *nkx2.2:GFP+* cells in the forebrain from 24 to 36 hpf were HuC/D+ (Figure 16 A,D,G). Throughout this time course, HuC/D, *nkx2.2:GFP* double positive cells were most prevalent in the telencephalon (Figure 16 B,C). Surprisingly, the earliest telencephalic neurons that were positioned about the presumptive AC region were largely all positive for *nkx2.2:GFP* (Figure 16). As the *nkx2.2:GFP+*population expanded in the telencephalon over time, this population occupied more dorsal-anterior positions, and the axonal projections of HuC/D+, *nkx2.2:GFP+* neurons were observed to contribute directly to the AC (Figure 16 D,G, upper bracket). Interestingly, at 28 and 36 hpf, the HuC/D+ cells more ventral to the AC were negative for *nkx2.2:GFP* expression (Figure 16 D,G, lower bracket). We also noted an expansion of *nkx2.2:GFP* expressing cells within the far lateral forebrain, however these cells were much deeper than, and farther from, the commissure as well as largely absent of HuC/D labeling (Figure 16 D,E,G,H yellow arrows). By 36 hpf, a small fraction of the neurons in the diencephalon were *nkx2.2:GFP*, HuC/D double positive (Figure 16 H-J). These results, taken together, suggest that *gfap* and *olig2* positive cells represent important stem cells in the forebrain, primarily for the generation of neuronal populations, particularly in the telencephalon (Figure 17).

**Figure 16:**
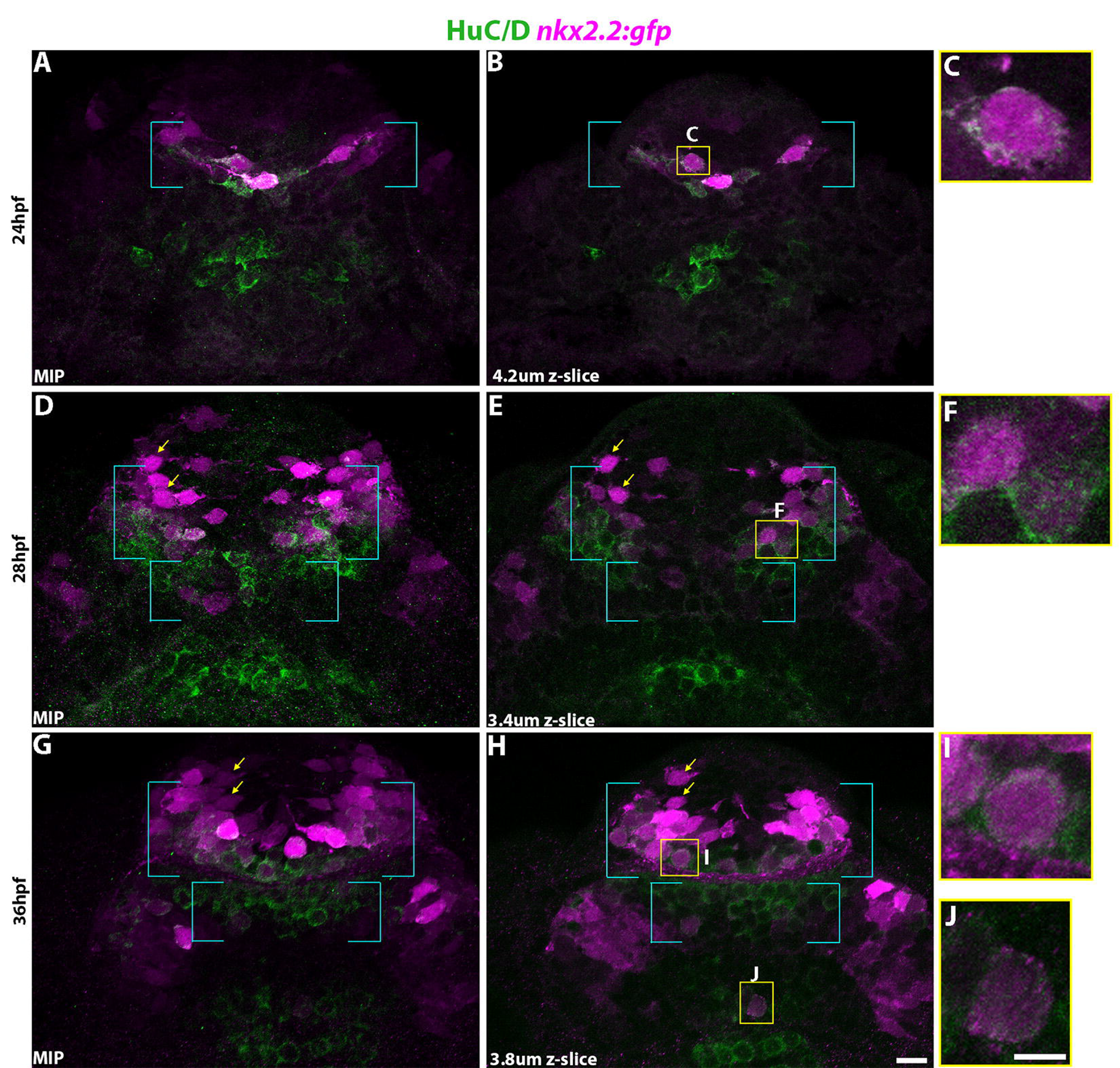
Despite expression of *tg(nxk2.2:gfp)* in the forebrain, resident oligodendrocyte progenitor cells have not adopted a myelinating fate prior to 36 hpf. A,D,G) Frontal view color composite MIPs and (B,E,H) coronal z-selection from (A,D,G) of *tg(nxk2.2:gfp)* (magenta) embryos labeled with anti-HuC/D (green) at 24 (A,B), 28 (D,E), and 36 hpf (G,H). C,F,I,J) Individual selected cell inset composites from (B,E,H; scale bar = 11*μ*m). Yellow arrows) (D,E,G,H) HuC/D negative *nxk2.2:gfp+* populations (scale bar = 4.7*μ*m). Upper teal bracket) population region of largely HuC/D+ *nxk2.2:gfp+* cells which contribute to the AC. Lower teal bracket) population region of largely HuC/D+ *nxk2.2:gfp-*cells. hpf, hours post fertilization

**Figure 17:**
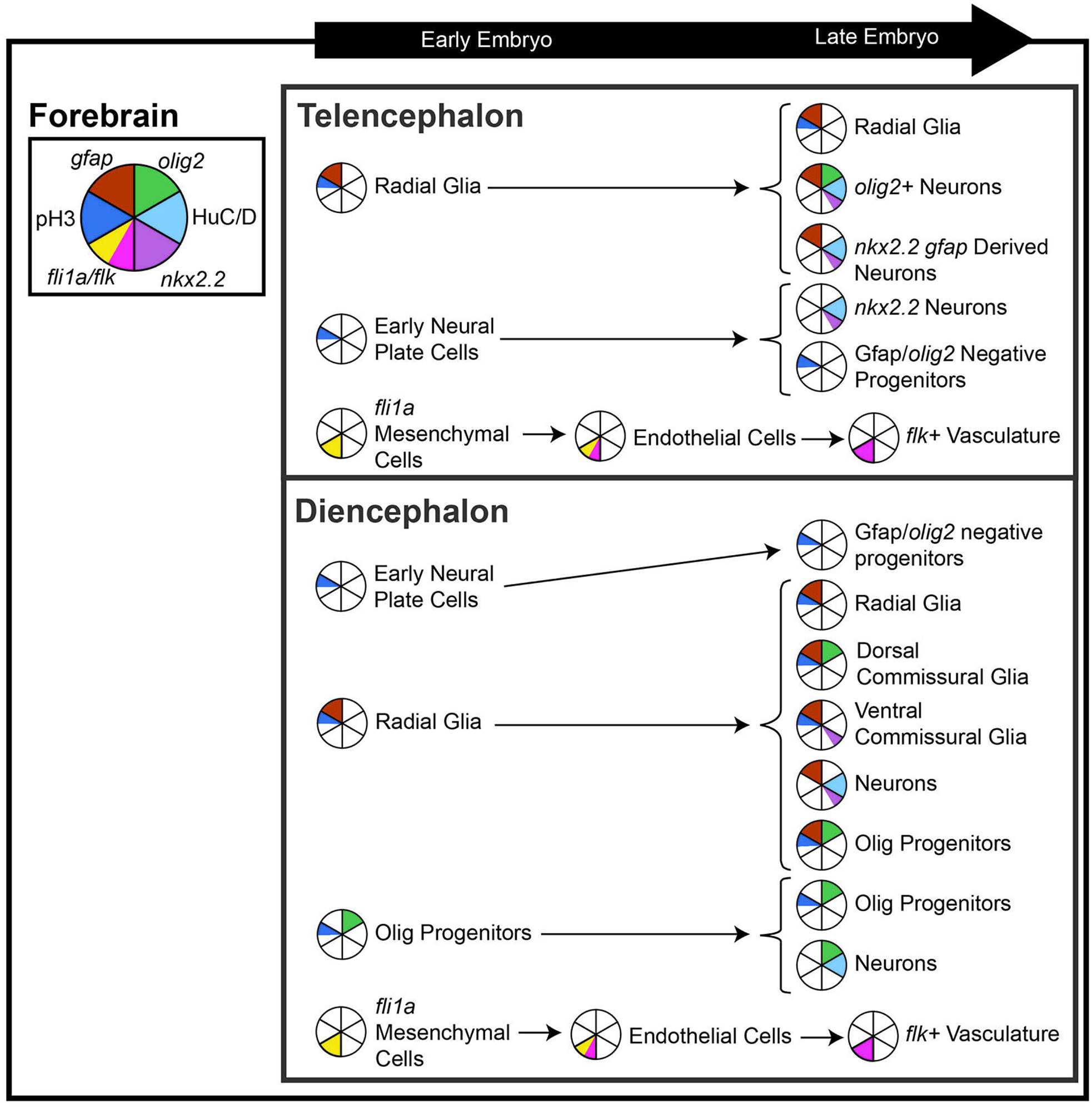
Progenitor populations and the ultimate fates of those cells are varied between the diencephalon and telencephalon, resulting in substantively different compositions despite close spatial proximity. Forebrain) The as yet unlabeled neuroepithelial cell and early infiltration of the region by vascular and crest derived cells as early as 12 hpf set the stage for early forebrain construction. Later, *fli1a/flk+* cells infiltrate the diencephalon and telencephalon and begins sprouting between 24 and 28 hpf. Anterior Embryo) By 20-22 hpf, *fli1a+* cells cover the anterior embryo. Telencephalon) *gfap+* cells and unlabeled cells are observed to give rise to further proliferative populations, and multiple lineages of neurons. Diencephalon) Earlier specification of *olig2+*populations results in an additional population of neuroprogenitors, negative for *gfap*, in addition to in addition to an additional proliferative population of *olig2+gfap+* cells and previously unobserved POC associated glial morphologies.

### 3.5 Vascular Cell and Commissural Axon Interactions

We previously observed the presence of *fli1a+* endothelial cells at 48 hpf in the forebrain in proximity to the developing commissures (Figure 1). How these vascular cells interact with and influence commissure development is unknown. We performed a more detailed time course of *tg(fli1a: GFP)* expression during the development of the AC and POC from 24 and 36 hpf. Interestingly, relatively superficial *fli1a:GFP+* cells moved en mass into the forebrain along bilateral paths of the presumptive anterior and post optic commissure regions (Figure 18, Sup. Mov. 10 A-C). Although, at 24 hpf we did observe a curious *fli1a:GFP+* cell extending a vascular-like process (Figure 18 A, yellow arrow), all other cells had a more mesenchymal morphology (Figure 18 A, red arrow). By 28 and 36 hpf, these *fli1a:GFP+* mesenchymal cells were overlapping both commissures of the forebrain (Figure 18 B,C). In the telencephalon, it was not until 36 hpf that we saw distinctive *fli1a:GFP+* cell morphology consistent with endothelial cell vasculogenesis (Lawson and Weinstein, 2002). More specifically, *fli1a:GFP+* endothelial-like cells approached the telencephalic midline with protruding cell processes from bilateral positions just ventral to the anterior commissure (Figure 18 C, yellow arrow).

**Figure 18:**
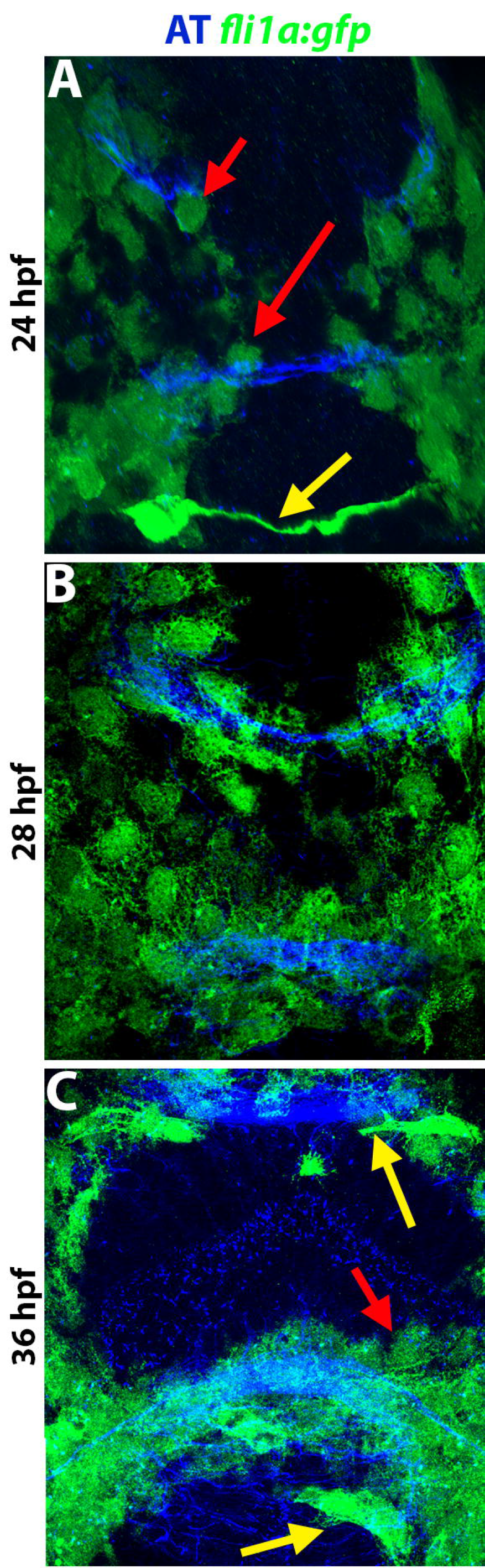
*fli1a+* cells infiltrate the zebrafish forebrain prior to 24 hpf, and display melanocytic and vascular morphologies at 24 hpf. A-C) Frontal color composite MIPs of *tg(fli1a:gfp)* (green) labeled with anti-AT (blue) of 24, 28, and 36 hpf embryos. Arrows) Cell populations with noted mesenchymal (red) or vascular (yellow) morphology. hpf, hours post fertilization

Between 24 hpf and 36 hpf where we had previously observed cells of vascular morphology in the *tg(fli1a)* line, *flk1:GFP+* vascular cells were seen similarly spanning across the telencephalic midline (Figure 19 A-C). Additionally, at 24 and 28 hpf *flk1:GFP+* cells were observed intruding into the forebrain region, ventral to the POC (Figure 19 A,G). By 28 hpf, some *flk1:GFP+* cells were observed entering the telencephalon in concert with the astroglial cells of the AC (Figure 19 G,H,I). Interestingly, we observed that *flk1:GFP+* cells adopted morphologies consistent with the tubular structure of vasculature as early as 28 hpf, and these cells remained in relative close proximity to both the AC (blue) and associated telencephalic glial bridge (Figure 19 S-U). This association with the telencephalic glial bridge continued through 36 hpf, where relative correspondence between AC axons was seen both lateral to, and at, the midline (Figure 19 G-I). Interestingly, these *flk1:GFP+* cells were not touching directly the AC but rather observed to be at a small and regular distance from the anterior commissure along its ventral side (Figure 19 H). No *flk1:GFP+* (or *fli1a:GFP+)* vascular cells were seen in the diencephalon at these times; instead, *flk1:GFP+* cells were situated both ventrally and superficially to the developing POC in the diencephalon (Figure 19 H,I,N,O). However, by 48 hpf, these diencephalic *flk1:GFP+*cells had migrated over the POC within Gfap+ astroglia and projected dorsally where they met up with *flk1:GFP+* vasculature in the telencephalon growing ventrally at the midline (Figure 19, Sup. Mov. 7; Sup. Mov. 11).

**Figure 19:**
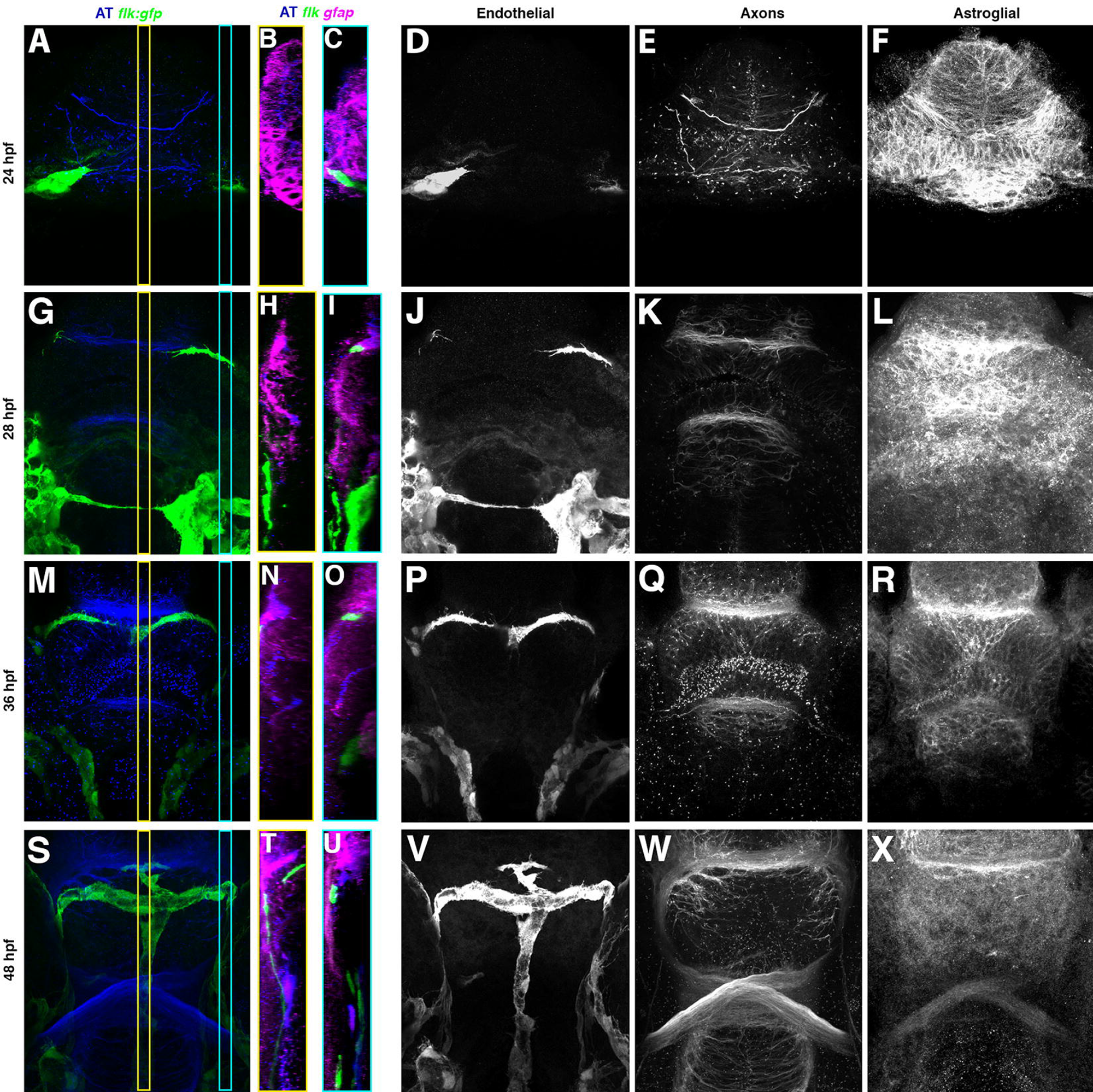
*flk1+* cells form ventrally oriented vascular in close proximity to the AC and interact with the telencephalic glial bridge. A,G,M,S) Color composite frontal MIPs of *tg(flk1:gfp)* (green) labeled with anti-Acetylated Tubulin (blue) of 24, 28, 36, and 48 hpf embryos with single channel monochromes of (A,G,M,S) showing *flk1:gfp* (D,J,P,V), anti-AT (E,K,Q,W) and *gfap:mCherry-caax* (F,L,R,X). B,H,N,T) Midline sagittal MIP of yellow outlined slice in (A,G,M,S) with added *gfap:mCherry-caax* (magenta) channel. C,I,O,U) Lateral and sagital MIP of cyan outlined slice in (A,G,M,S) with added *gfap:mCherry-caax* (magenta) channel. hpf, hours post fertilization

## 4 Discussion

Our study sought to characterize the myriad of cells that interact with commissural axons during the development of the embryonic forebrain. Historically, most studies of the developing brain have focused on the behavior of axons themselves or perhaps the influence that one cell type may have on the developing axonal anatomy (Kaprielian, Runko, and Imondi, 2001; Zou and Lyuksyutova, 2007; Fothergill et al., 2014; Tosa et al., 2015; Friocourt and Chédotal, 2017; Bayraktar et al., 2015). Such focused approaches fail to incorporate the potential contributions that an array of cooperating cell populations and their complex interactions may have in the guidance mechanisms underlying commissure development. To identify which cell populations may be directly influencing commissure formation in the forebrain, we attempted to simultaneously assay the spatial and temporal development of different progenitor populations, glial cells, neurons, and endothelial cells with pathfinding commissural axons. This comprehensive characterization has led to a better understanding of the changes that occur to the cellular architecture associated with the commissural regions of the telencephalon and diencephalon over time. We summarized the findings of this characterization in a visual model (Figure 20, Sup. Mov. 12 A-C).

**Figure 20:**
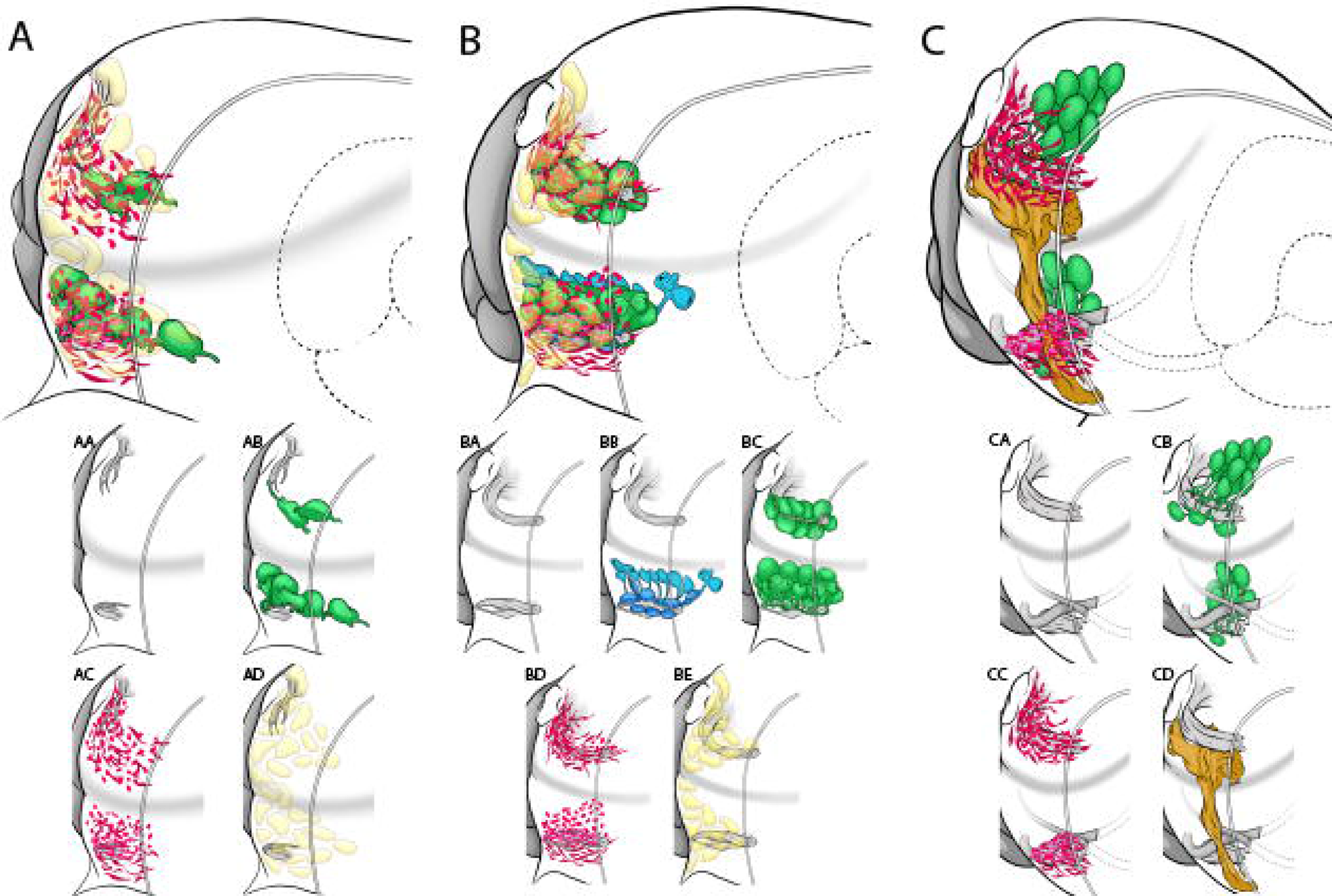
An illustrated model of the commissural cell architecture of the zebrafish forebrain. A-C) Model of the observed positions and cell populations extant at 22, 28 and 48 hpf respectively. The illustration shows different cell populations relative to the telencephalon (above shaded line) and diencephalon (below shaded line) from a partial frontal and lateral view that has a transverse sectional view through it (double lines). The eye and optic nerve on the cut side is shown as a dotted line for reference. Each cell population is show relative to commissures such that axons of the POC and AC are show at each time point (grey), *gfap+* astroglia (blue), *olig2+* progenitor cells (green), ZRF1-4 immunolabeling (red), and fli1a+ cells and vasculature (cream and brown respectively). AA-CD) shows each cell population separately for clarity of that populations architecture alone. Distinct radial glial morphologies are only depicted at the 28 hpf stage because this the only time point for which our chimeric analysis took place.

### 4.1 Cell heterogeneity in the developing embryonic forebrain

Just prior to the exploration of the midline by pathfinding axons, the presumptive commissural regions are prefigured by several cell growth substrates. Differential labeling of Gfap and Zebrafish radial fiber antibodies define condensed bridge-like swaths of astroglial cells spanning the telencephalic and diencephalic midlines. We discovered a remarkable correspondence of *fli1a+* superficial mesenchymal cells specifically with both commissural regions before and during early axon pathfinding across the midline. Moreover, *gfap+* and *olig2+* progenitor cell populations were present throughout commissure development, though timing and distribution differences were found between the telencephalon and diencephalon. In the diencephalon, three different astroglial cell morphologies were observed, two positioned at the dorsal and ventral sides of the commissure as well as ventricular zone radial glia with endfeet terminating directly upon the POC. In addition, *olig2+* progenitor cells (OPCs) occupied dorsal restricted positions relative to the POC, whereas they dynamically surrounded the AC in the telencephalon. These progenitor cells were seen to exhibit proliferative markers during the period of commissure formation. Significant portions of the telencephalic OPCs were derived from Gfap+ stem cells, whereas the origin of OPCs in the diencephalon appeared more diversified. Lastly, we showed early development of telencephalic blood vessels running parallel to, yet subtly separated from, the anterior commissure, as well as distinct dorsally growing endothelial sprouts along the midline of both the telencephalon and diencephalon (Figure 20, Sup. Mov 12 A-C). Our model suggests that a diverse cellular architecture exists to setup where commissures will form, which establishes axonal structures that then serve as a scaffold to further reinforce the development of neuronal, glial and endothelial cell differentiation. Additionally, we have revealed a significant diversity in the types of cells that appear to directly interact with pathfinding commissural axons during forebrain development. First, we noted the early presence of *fli1a-GFP+* cells that appear to exhibit mesenchymal morphology and, most remarkably, follow the track of the forebrain commissures both before and during the times when axons are crossing the midline (Figure 18). These initial stages of commissure development are known to be too early for vasculogenesis (Lawson and Weinstein, 2002). Interestingly, *tg(fli1a-GFP)*transgene expression has been documented in cranial neural crest cells particularly during pharyngeal arch morphogenesis and subsequent cartilage formation during craniofacial development (Kimmel, Miller, and Keynes, 2001; Schilling and Kimmel, 1994; Graham, Begbie, and McGonnell, 2004; Trainor and Tam, 1995; Kaucka et al., 2016; Bohnsack, Gallina, and Kahana, 2011). We propose that these *fli1a-GFP+* cells in the forebrain may constitute some of the most anterior streams of cranial neural crest cells that are potentially navigating through the presumptive commissure regions. This is a particularly interesting possibility because cranial neural crest cells have recently been implicated in secreting various signals during their migration through the zebrafish eye to influence the extracellular matrix necessary for optic stalk development as well as through the avian somite to regulate myogenesis (Bryan et al., 2020; Rios et al., 2011). These *fli1a-GFP+* cells are present at the critical times and with the spatial refinement to directly influence the guidance substrate for commissural axons. Future investigations are therefore merited to directly test whether these *fli1a-GFP+* cells are in fact of neural crest origin and necessary for the proper guidance of commissural and retinal ganglion cell axons as well as for neuronal and glial cell differentiation in the forebrain.

Our characterization of progenitor and glial cell populations in the embryonic forebrain also demonstrated a surprising level of early cell heterogeneity affiliated with the commissure regions. The commissure-prefiguring and bridge-like distribution of Gfap labeling has long suggested their involvement with the guidance of axons across the midline (Marcus and Stephen S Easter, 1995; Barresi et al., 2005; Schwartz et al., 2020). In addition, we demonstrated here that at least in the diencephalon, Gfap labeling demarcates a portion of the glial bridge, while labeling with Zrf2-4 revealed more expansive and Gfap overlapping domains for the diencephalic glial bridge (Figure 5, Figure 6). Furthermore, we discovered the presence of three different *gfap+* astroglial morphologies, which appear to be strategically positioned in contact with the POC (Figure 10, Figure 9). Interestingly, despite the conventional premise that radial glial cells are the only astroglial cell type in zebrafish, a very recent exploration of the adult zebrafish olfactory bulb revealed a unique astroglial morphology divergent from the characteristic radial cell morphology of radial glia (Scheib and Byrd-Jacobs, 2020). We propose that these divergent glial morphologies may arise as early as the embryonic brain, and persist throughout larval and juvenile development and into the adult. For such a hypothesis to be tested would require long-term cell lineage analyses as well as the identification of more robust astroglial type markers in zebrafish. Determination of the antigenicity of the Zebrafish radial fiber 2-4 antibodies could represent a first step toward the identification of new astroglial markers.

### 4.2 Developmental heterochrony between the telencephalon and diencephalon

By conducting our analysis over several key time periods during commissure formation, we cataloged clear differences in the developmental timing of similar cell types between the diencephalon and telencephalon. The heterochrony of the forebrain commissures has certainly been well documented, such that the diencephalic midline is first pioneered by POC axons at around 23 hpf, which is then followed by the AC at 24.5hpf and finally the optic chiasm at 36hpf (Figure 2); (Hjorth and Key, 2004; Wilson, Ross, et al., 1990; Wilson and Easter, 1991). Interestingly, despite the earlier commissure formation in the diencephalon, we show a significantly later differentiation of *HuC+* neurons in the diencephalon as compared to neuronal differentiation in the telencephalon (Figure 14). The POC is known to be pioneered by bilaterally positioned neuronal populations known as the ventral rostral clusters that reside much deeper inside the diencephalon; in contrast, our labeling of telencephalic *HuC+* neurons showed that they were directly contributing to the AC (Figure 14). This heterochronic neuronal differentiation is further supported by the earlier *nkx2.2* driven transgene expression in neurons of the 24 hpf telencephalon as compared to the first *nkx2.2+* neurons appearing in the 36hpf diencephalon. We find this to be a perplexing timing difference when considering the expansive and diverse progenitor populations that exist in the diencephalon (Figure 12). This comparatively delayed differentiation in the diencephalon is further reinforced by our characterization of vasculogenesis, for which clear blood vessels are forming in the 30hpf telencephalon and not in the diencephalon until 48hpf (Figure 19). These results taken together suggest that different stem and progenitor cell programs exist in the telencephalon and diencephalon that result in these differences in the timing of cell differentiation. We speculate that one reason why the diencephalon may need to maintain a less differentiated state for a longer period of time is to keep in place an overall guidance environment suitable for the later crossing of retinal ganglion cell axons coming from the eyes to form the optic chiasm.

### 4.3 The role of transient cell fates in building the brain

It is a common tendency for researchers to focus more heavily on the beginning and ending periods of a cell’s maturation; however, the period of embryonic development represents but a transient moment in the life of the cells actively building the embryo. This investigation has highlighted the important roles that transient progenitor cells may play in the building of the brain. Through our characterization of *olig2* and *gfap* driven transgene expression we were able to discern lineage, position, and morphology differences with the progenitor cell populations of the forebrain. OPCs were significantly different between the telencephalon and diencephalon with respect to their architecture around the AC and POC, respectively. OPCs come to occupy both dorsal and ventral positions around the AC, while OPCs almost exclusively reside on just the dorsal portions of the POC (Figure 11). Moreover, while most of the telencephalic OPCs co-expressed gfap-driven transgene expression, many of the more medially positioned OPCs in the diencephalon did not (Figure 12). Both of these astroglial (*gfap+)* and OPC populations were proliferative throughout the early stages of commissure development (Figure 12). Moreover, we showed that these progenitor cells did contribute, in some portion, to the neuronal differentiation of Serotonergic neurons, rather than MBP+ oligodendroglia (Figure 13, Figure 15). Interestingly though, during a majority of the period of time the forebrain commissures were forming, these progenitor cell populations were also extending cell processes that were directly interacting with pathfinding axons (Figure 11). This suggests that these transient progenitor populations of astroglia and OPCs serve in part as the axon growth substrate for both the POC and AC. It is likewise plausible, that as commissures are pioneered they also serve to reinforce the further recruitment of progenitor cells types at the commissure’s position. Moreover, as mentioned above, we hypothesize that migrating cranial neural crest cells (*fli1a:GFP+*) stream through the presumptive commissural regions, for which these highly transient cells may also play an active role in contributing to the axon substrate milieu. A similar role has been suggested for the migrating population of neural progenitor cells that constitutes the “glial sling,” which are cells required for optic chiasm development in the mouse (Silver et al., 1982; Shu, Puche, and Richards, 2003). Therefore, it is important to understand not just the starting and ending stages of a tissue’s maturation but the complex interactions with transient cell types that may occur only momentarily during its construction.

### 4.4 Architecture should inform guidance

One of core fundamental concepts in biology is that “form fits function,” and in this study we describe the cellular building blocks of the developing forebrain, and thereby should inform the functional units at play. As we have grounded this investigation on commissure formation, we looked to the cellular structures present at different times to glean insight upon the types of cell guidance mechanisms that may be functioning to position the cells and axons of the forebrain correctly. The coincident localization of astroglial cells and OPCs with pathfinding commissural axons, suggests these progenitor cell types are directly functioning as a growth substrate for these axons and/or regulating their guidance. We have shown previously that Gfap+ astroglia express *slit1a*, which is a member of the slit family of guidance cues most notably regarded for their repellent functions (Barresi et al., 2005; Tosa et al., 2015; Fothergill et al., 2014; Kaprielian, Runko, and Imondi, 2001; Kidd, Bland, and Goodman, 1999; Brose et al., 1999). However, the overlapping position of these astroglial cells with AC and POC axons would suggest these cells provide a more permissive axon guidance function. We recently demonstrated that Slit1a may function rather to first guide the condensation of astroglial cells into prefigured bridges that then supports midline crossing of commissural axons (Schwartz et al., 2020). Our analysis here further breaks down the varied parts of the astroglial bridge into three different morphologies and three different Zrf defining domains. How each of these cells and their respective cell processes function to influence commissural axon guidance remains unknown. It is nevertheless intriguing to speculate about the permissive guidance role that the elaborate endfeet of radial glial cells play as they extend across the superficial-most portions of commissures. Furthermore, it remains unknown whether dorsal versus ventral commissural astroglial cells differ in their axon guidance roles. While both populations are *gfap+*, whether they arise from the same wave of progenitor cell differentiation, and equally contribute to guidance, remains to be discovered, as does the determination of their ultimate lineage fates. Nevertheless, the differential morphology and positioning of these two populations of glial, and the differences in their associations with POC axons, highlights the need for further investigation of whether these cells differentially contribute to commissural axon guidance or forebrain glial populations.

In comparison to the astroglial cell populations, the role OPCs play in axon guidance is less predictable. Although, we demonstrated that OPC in the telencephalon are directly generating neurons that pathfind along the AC, OPCs are also interlacing multiple cell processes into the developing commissures (Figure 8, Figure 11). We did not observe any *nkx2.2+* OPCs in proximity to the POC or AC that might suggest an oligodendroglia fate and consequent myelination function, therefore the current role of these OPC-commissure interactions remain a mystery. Of particular fascination was the gradual coalescence of OPCs to the dorsal midline of the POC, which corresponded uniquely to the positioning of the optic chiasm. This characterization of OPC development with commissure formation highlights a new area of contact-mediated guidance yet to be explored.

Lastly, the progressive elaboration of blood vessels first in the telencephalon and then in the diencephalon fall along conserved dimensions that suggest the reoccurring involvement of stable guidance systems. For instance, in the telencephalon, endothelial cells first grow bilaterally toward the midline parallel to but displaced ventrally from the already established AC (Figure 19). How is this exacting distance from the AC maintained during endothelial cell migration and vasculogenesis? Furthermore, once at the midline these endothelial cells of the telencephalon dramatically alter this migratory direction and dive deeper to grow along the midline yet now behind the AC (Figure 19). In contrast, endothelial cells in the diencephalon are restricted to growing along the superficial midline over the POC and optic chiasm (Figure 19). These types of cell behaviors are characteristic responses to environmental guidance cues (Fujiwara, Ghazizadeh, and Kawanami, 2006; London and Li, 2011; Adams and Eichmann, 2010), and the next step investigations will be to define which cellular and molecular cues of the forebrain are responsible for these endothelial cell migratory choices. By embarking on a more comprehensive analysis of the diverse array of cell interactions that occur during commissure formation in the zebrafish forebrain, we have not surprisingly discovered an equally diverse array of new questions to solve.

## Data Sharing

The data that support the findings of this study are available from the corresponding author upon reasonable request.

## Acknowledgments

We would like to thank Alicia Famiglietti, Cassie Kemmler, Narendra Pathak, Risha Sinha, Rachael Stein, Carla Valez, and Paula Zaman for all their supportive contributions during the course of this research. We would also like to thank the Smith College Center for Microscopy and its manager, Judith Wopereis, and the Smith Animal Care facility for their constant support and technical assistance throughout this work. Lastly, we are extremely appreciative of all constructive discourse provided by the entire Barresi lab from the first experiments to the final manuscript. This research was generously supported by the National Science Foundation [IOS-1656310] and by Smith College for both undergraduate and graduate student support.

## Conflicts of Interest

The authors declare no conflicts of interest.

## Supplementary Information

**Supplementary Movie 1A: Gfap forms glial bridges which are interlaced through the forebrain commissures.** 3D rendering of confocal z-stack of a 48 hpf embryo labeled with anti-AT (blue) and anti-Gfap (green). Left panel rotates counterclockwise around the anterior-posterior vertical axis, starting from a ventral perspective. The right panel shows the same sample rotating counterclockwise around the anterior-posterior horizontal axis, starting from a lateral prospective and moving towards a ventral prospective.

**Supplementary Movie 1B: *gfap+* glial cell membranes overlap with axons of forebrain commissures.** 3D rendering of confocal z-stack of a 48 hpf *tg(gfap:gfp-caax)* (green) embryo labeled with anti-AT (blue). Left panel rotates counterclockwise around the anterior-posterior vertical axis, starting from a dorsal perspective. The right panel shows the same sample rotating counterclockwise around the anterior-posterior horizontal axis, starting from a lateral prospective and moving towards a ventral prospective.

**Supplementary Movie 1C: *Olig2+* cells densely populate the telencephalon and extend processes towards the AC.** 3D rendering of confocal z-stack of a 48 hpf *tg(olig2:gfp)* (green) embryo labeled with anti-AT (blue). Left panel rotates clockwise around the anterior-posterior vertical axis, starting from a ventral perspective. The right panel shows the same sample rotating counterclockwise around the anterior-posterior horizontal axis, starting from a lateral prospective and moving towards a dorsal prospective.

**Supplementary Movie 1D: *fli1a+* cells track the tract of the AC to form vasculature in the telencephalon.** 3D rendering of confocal z-stack of a 48 hpf *tg(fli1a:gfp)* (green) embryo labeled with anti-AT (blue). Left panel rotates clockwise around the anterior-posterior vertical axis, starting from a ventral perspective. The right panel shows the same sample rotating clockwise around the anterior-posterior horizontal axis, starting from a lateral prospective and moving towards a dorsal prospective.

**Supplementary Movie 2A: Glial membranes and Gfap form overlapping but noncoincident glial bridges at 22 hpf, prior to commissure formation.** Transverse slices of 22 hpf *tg(gfap:cherry-caax)* (magenta) and immunolabeled with anti-AT (blue) and anti-Gfap (green). Video begins and ends laterally at each rostral cluster and passes through the midline. Anterior is to the left.

**Supplementary Movie 2B: The Gfap+ glial bridge is superficial to the forebrain commissures, while glial membranes are instead coincident with POC axons at 24 hpf.** Transverse slices of 24 hpf *tg(gfap:cherry-caax)* (magenta) and immunolabeled with anti-AT (blue) and anti-Gfap (green). Video begins and ends laterally at each rostral cluster and passes through the midline. Anterior is to the left.

**Supplementary Movie 2C: The Gfap+ glial bridge is superficial to the forebrain commissures, while glial membranes are densely coincident with POC axons at 28 hpf.** Transverse slices of 28 hpf *tg(gfap:cherry-caax)* (magenta) and immunolabeled with anti-AT (blue) and anti-Gfap (green). Video begins and ends laterally at each rostral cluster and passes through the midline. Anterior is to the left.

**Supplementary Movie 2D: The Gfap+ glial bridge is superficial to the forebrain commissures, while glial membranes are instead coincident with POC axons at 36 hpf.** Transverse slices of 36 hpf *tg(gfap:cherry-caax)* (magenta) and immunolabeled with anti-AT (blue) and anti-Gfap (green). Video begins and ends laterally at each rostral cluster and passes through the midline. Anterior is to the left.

**Supplementary Movie 3: Timelapse microscopy showing first pathfinding axons of the POC and AC.** Dorsal view of forebrain of *tg(gap43:gfp)* (green) embryo from 22 hpf to 26 hpf.

**Supplementary Movie 4: Pioneering axons of the AC navigate in association with and through *gfap+* glial membranes.** Dorsal view of forebrain of *tg(gap43:gfp)* (red) and *tg(gfap:cherry-caax)* (green) embryo from 22 hpf to 30 hpf with 5 min time intervals.

**Supplementary Movie 5A: 3D visualization of *gfap+* cells with radial glial morphology which surround the POC.** Transplanted population of cells at 28 hpf from *tg(gfap:nls-mcherry)* (red) and *tg(gfap:gfp-caax)* (green) embryo labeled with anti-AT (blue). Axes approximately correspond to x = lateral, y = dorsal, z = anterior.

**Supplementary Movie 5B: 3D visualization of *gfap+* cells with dorsal commissural glial morphology are found dorsal to the POC.** Transplanted population of cells at 28 hpf from *tg(gfap:nls-mcherry)* (red) and *tg(gfap:gfp-caax)* (green) embryo labeled with anti-AT (blue). Axes approximately correspond to x = lateral, y = dorsal, z = anterior.

**Supplementary Movie 5C: 3D visualization of *gfap+* cells with ventral commissural glial morphology are found ventral to POC.** Transplanted population of cells at 28 hpf from *tg(gfap:nls-mcherry)* (red) and *tg(gfap:gfp-caax)* (green) embryo labeled with anti-AT (blue). Axes approximately correspond to x = lateral, y = dorsal, z = anterior.

**Supplementary Movie 5D: 3D rendering of *gfap+* cells with radial glial morphology.** 3D reconstruction and 360 degree rotational rendering of isolated cell with radial glial morphology.

**Supplementary Movie 5E: 3D rendering of *gfap+* cells with dorsal commissural glial morphology are found dorsal to the POC.** 3D reconstruction and 360 degree rotational rendering of isolated cell with dorsal commissural glial morphology.

**Supplementary Movie 5F: 3D rendering of *gfap+* cells with ventral commissural glial morphology.** 3D reconstruction and 360 degree rotational rendering of isolated cell with ventral commissural glial morphology.

**Supplementary Movie 6A: *olig2+* cells heavily populate the diencephalon and lightly populate the telencephalon at 22 hpf.** 3D rendering of *tg(olig2:gfp)* (green) forebrain immunolabeled with anti-AT (blue). Left panel begins from a frontal prospective and rotates clockwise 360° around the anterior-posterior axis. The right panel is the same sample and begins from a more dorsal prospective and rotates in phase with the left panel. AT signal intensity is increased for structural visualization purposes.

**Supplementary Movie 6B: *olig2+* cells heavily populate the diencephalon and increase in cell number in the telencephalon at 24 hpf.** 3D rendering of *tg(olig2:gfp)* (green) forebrain immunolabeled with anti-AT (blue). Left panel begins from a frontal prospective and rotates clockwise 360° around the anterior-posterior axis. The right panel is the same sample and begins from a more dorsal prospective and rotates in phase with the left panel.

**Supplementary Movie 6C: The *olig2+* population has greatly increased in the telencephalon by 28 hpf.** 3D rendering of *tg(olig2:gfp)* (green) forebrain immunolabeled with anti-AT (blue). Left panel begins from a frontal prospective and rotates clockwise 270° around the anterior-posterior axis. The right panel is the same sample and begins from a more dorsal prospective and rotates in phase with the left panel.

**Supplementary Movie 6D: *olig2+* populations have greatly expanded by 36 hpf and exhibit many multiprocess contacts with the forebrain commissures.** 3D rendering of *tg(olig2:gfp)* (green) forebrain immunolabeled with anti-AT (blue). Left panel begins from a frontal prospective and rotates clockwise 270° around the anterior-posterior axis. The right panel is the same sample and begins from a more dorsal prospective and rotates in phase with the left panel.

**Supplementary Movie 7A: *gfap+* and *olig2+* populations are forebrain progenitors at 22 hpf.** Scrolling through frontal z-stack of a 22 hpf *tg(*gfap:nls-mcherry (magenta); olig2:gfp (green) embryo labeled for pH3 (blue). Movie begins at the most superficial/anterior slice and scrolls to deeper/more posterior slices, spanning the entire commissural region.

**Supplementary Movie 7B: *gfap+* and *olig2+* populations are each discrete forebrain progenitors at 24 hpf.** Scrolling through frontal z-stack of a 24 hpf *tg(*gfap:nls-mcherry (magenta); olig2:gfp (green) embryo labeled for pH3 (blue). Movie begins at the most superficial/anterior slice and scrolls to deeper/more posterior slices, spanning the entire commissural region.

**Supplementary Movie 7C: Telencephalic and diencephalic *gfap+* and *olig2+* differentially contribute to glial lineages.** Scrolling through a frontal z-stack of a 28 hpf *tg(*gfap:nls-mcherry) (magenta) olig2:gfp (green) double transgenic embryo labeled for pH3 (blue). Movie begins at the most superficial/anterior slice and scrolls to deeper/more posterior slices, spanning the entire commissural region.

**Supplementary Movie 8A: *nxk2.2:gfp+* forebrain populations are predominantly found in the telencephalon, and are not coincident with *olig2:mcherry+* populations at 28 hpf.** 3D rendering of a 28 hpf *tg(nxk2.2:gfp)* (green), *tg(olig2:mcherry)* (purple) forebrain labeled with anti-AT (blue). Left panel begins from a frontal prospective and rotates counterclockwise 270° around the anterior-posterior axis. The right panel is the same sample and begins from a more dorsal prospective and rotates in phase with the left panel.

**Supplementary Movie 8B: *nxk2.2:gfp+* telencephalic populations have greatly expanded, but are still *olig2:mcherry* - at 36 hpf.** 3D rendering of a 36 hpf *tg(nxk2.2:gfp)* (green), *tg(olig2:mcherry)* (purple) forebrain labeled with anti-AT (blue). Left panel begins from a frontal prospective and rotates clockwise 300° around the anterior-posterior axis. The right panel is the same sample and begins from a more dorsal prospective and rotates in phase with the left panel.

**Supplementary Movie 9A: *gfap+* and *olig2+* lineage neurons are evident at 24 hpf.** Scrolling through frontal z-stack of a 24hpf *tg(gfap:nls-mcherry* (magenta); *olig2:gfp* (green)) forebrain labeled with anti-HuC/D (blue). Movie begins at the most superficial/anterior slice and scrolls to deeper/more posterior slices, spanning the entire commissural region.

**Supplementary Movie 9B: *gfap+* and *olig2+* lineage neurons have increased in number at 28 hpf, but differentially contribute to telencephalic versus diencephalic populations.** Scrolling through frontal z-stack of a 28hpf*tg(gfap:nls-mcherry)* (magenta); *olig2:gfp* (green)) forebrain labeled with anti-HuC/D (blue). Movie begins at the most superficial/anterior slice and scrolls to deeper/more posterior slices, spanning the entire commissural region.

**Supplementary Movie 10A: *fli1a+* cells form vascular like processes ventral and superficial to the POC at 24 hpf.** 3D rendering of a 24hpf *tg(fli1a:gfp)* (green) forebrain immunolabeled with anti-AT (blue). Left panel begins from a frontal prospective and rotates counterclockwise 300° around the anterior-posterior axis. The right panel is the same sample and begins from a more dorsal prospective and rotates in phase with the left panel.

**Supplementary Movie 10B: *fli1a+* cells follow the tract of the AC as they form telencephalic projections at 28 hpf.** 3D rendering of a 28hpf *tg(fli1a:gfp)* (green) forebrain immunolabeled with anti-AT (blue). Left panel begins from a frontal prospective and rotates clockwise 300° around the anterior-posterior axis. The right panel is the same sample and begins from a more dorsal prospective and rotates in phase with the left panel.

**Supplementary Movie 10C: *fli1a+* cells form vascular-like structures in the forebrain, while superficially coating the forebrain at 36 hpf.** 3D rendering of a 36hpf *tg(fli1a:gfp)* (green) forebrain immunolabeled with anti-AT (blue). Left panel begins from a frontal prospective and rotates clockwise 300 around the anterior-posterior axis. The right panel is the same sample and begins from a more dorsal prospective and rotates in phase with the left panel.

**Supplementary Movie 11: *flk+* cells form vasular structures throughout the forebrain, with midline structures connecting the diencephalon and telencephalon.** 3D rendering of a 48 hpf *tg(flk1:gfp* (green); *gfap:cherry-caax* (magenta)) forebrain immunolabeled with anti-AT (blue). Left panel begins from a frontal prospective and rotates clockwise 360° around the anterior-posterior axis. The right panel is the same sample and begins from a more dorsal prospective and rotates in phase with the left panel.

**Supplementary Movie 12a: The early, 22 hpf, forebrain commissural cell architecture is constructed from several interacting cell types.** Movie of 22 hpf model starting with all cells then showing each cell type observed relative to the forebrain commissures individually, in the order of: forebrain, axons, *gfap+* glia, *olig2+cells*, Zrfs, *fli1a+* cells, and all channels combined. After individual cell models, channels are added to the total model in the order they appeared.

**Supplementary Movie 12b: The 28 hpf forebrain commissural cell architecture is exhibits heterochronous cell contributions between the telencephalon and diencephalon.** Movie of 28 hpf model showing each cell type observed relative to the forebrain commissures individually, in the order of: forebrain, axons, *olig2+* cells, *gfap+* glia, Zrfs, *fli1a+* cells, and all channels combined. After individual cell models, channels are added to the total model in the order they appeared.

**Supplementary Movie 12c: The 48 hpf forebrain commissural cell architecture exhibits many multiprocess cell types which interact with the forebrain commissures** Movie of 48 hpf model showing each cell type observed relative to the forebrain commissures individually, in the order of: forebrain, axons, *olig2+* cells, *flk+* cells, Zrfs, and all channels combined. After individual cell models, channels are added to the total model in the order they appeared.

## Notes

https://drive.google.com/drive/folders/1Vr9Dacy_BmFrWd5AkAC-DWtlooVB4TyD?usp=sharing

### Competing Interest Statement

The authors have declared no competing interest.

